# Splicing variation of BMP2K balances endocytosis, COPII trafficking and autophagy in erythroid cells

**DOI:** 10.1101/2020.05.05.079970

**Authors:** Jaroslaw Cendrowski, Marta Kaczmarek, Katarzyna Kuzmicz-Kowalska, Michal Mazur, Kamil Jastrzebski, Marta Brewinska-Olchowik, Agata Kominek, Katarzyna Piwocka, Marta Miaczynska

## Abstract

Intracellular transport undergoes remodeling upon cell differentiation, which involves cell type-specific regulators. Bone morphogenetic protein 2-inducible kinase (BMP2K) has been potentially implicated in endocytosis and cell differentiation but its molecular functions remained unknown. We discovered that its longer (L) and shorter (S) splicing variants regulate erythroid differentiation in a manner unexplainable by their involvement in AP-2 adaptor phosphorylation and endocytosis. However, both variants interacted with SEC16A whose silencing in K562 erythroid leukemia cells affected generation of COPII assemblies and induced autophagic degradation. Variant-specific depletion approach showed that BMP2K isoforms constitute a BMP2K-L/S regulatory system. Therein, L promotes while S restricts recruitment of SEC31A to SEC24B-containing COPII structures forming at SEC16A-positive ER exit sites. Finally, we found L to promote and S to restrict autophagic degradation. Hence, we propose that BMP2K-L favors SEC16A-dependent intracellular processes important for erythroid maturation, such as COPII trafficking and autophagy, in a manner inhibited by BMP2K-S.

## INTRODUCTION

In eukaryotic cells, vesicular transport underlies endocytosis and exocytosis, as well as it ensures the proper distribution of transmembrane proteins between different cellular compartments. The endocytic system delivers the transported cargo towards lysosomal degradation or recycling back to the plasma membrane (PM). Thus, endosomes often serve as signaling platforms for internalized receptors (Cendrowski et al, 2016; Irannejad et al, 2015). In turn, the exocytic machinery serves as a biosynthetic pathway for newly translated secretory or PM proteins. Although traditionally regarded as antagonistic, the endocytic and exocytic systems are now known to be widely interconnected as many proteins shuttle between endosomal and secretory compartments (Progida & Bakke, 2016).

Membrane trafficking involves vesicles which, depending on their origin, can be formed by one of protein coat assemblies (clathrin, COPI and COPII) (Gomez-Navarro & Miller, 2016). Clathrin assembles into lattices shaping vesicles transporting cargo from the PM or the *trans*-Golgi network (TGN) to the endolysosomal system (Robinson, 2015). Cargo incorporated into clathrin-coated vesicles is recognized by heterotetrameric adaptor protein (AP) complexes that define specific membrane donor sites. While AP-1 and AP-3 function in different sorting routes between the TGN and the endolysosomal pathway, the AP-2 complex acts in clathrin-mediated endocytosis (CME) (Park & Guo, 2014). AP-2 is a heterotetrameric complex consisting of two large (α and β2), one medium (μ2) and one small (σ2) subunits. COPI and COPII are involved in transporting cargo within the early secretory pathway which consists of the endoplasmic reticulum (ER), ER-Golgi intermediate compartment (ERGIC) and the Golgi stack (Wiseman et al, 2007). COPII forms vesicles that bud at the ER exit sites (ERES) and deliver cargo to the ERGIC and the Golgi (Venditti et al, 2014). COPI-coated vesicles transport cargo from the Golgi and the ERGIC back to the ER and between the Golgi cisternae (Arakel & Schwappach, 2018).

A feature that distinguishes COPII-coated carriers from other types of vesicles is that once they are formed, they immediately reach their target compartment, ERGIC located in close proximity to ERES (Lord et al, 2013). As initially characterized in yeast, COPII–mediated transport involves sequential recruitment of COPII complex components, including the Sar1 GTPase, the Sec23/Sec24 subcomplex, and the Sec13/Sec31 subcomplex. The Sec23/Sec24 inner shell sorts cargo into ER-derived vesicles while Sec13 and Sec31 polymerize forming their outer cage. The scission of budding vesicles occurs due to Sar1 GTPase activity which is stimulated by the assembled coat, particularly by the recruitment of Sec31 (Bielli et al, 2005; Lee et al, 2005; Sato & Nakano, 2005; Townley et al, 2008). COPII vesicle production is regulated by large protein Sec16 by two distinct mechanisms. The first one is a scaffolding function in organizing COPII assembly, confirmed for yeast and *Drosophila* Sec16 as well as mammalian SEC16A (Bhattacharyya & Glick, 2007; Connerly et al, 2005; Ivan et al, 2008; Martinez-Menarguez et al, 1999; Watson et al, 2006). The second role, shown for yeast Sec16, is to negatively regulate COPII turnover by inhibiting the Sec31 recruitment (Bharucha et al, 2013; Kung et al, 2012; Yorimitsu & Sato, 2012).

The vesicular trafficking pathways are regulated upon conditions that induce macroautophagy (hereafter referred to as autophagy) and contribute to autophagosome formation, maturation or autolysosomal clearance. Autophagosome biogenesis requires a contribution from both endocytic and secretory compartments (Lamb & Tooze, 2016). It is believed to be initiated at the ER and occur via incorporation of vesicles derived from ERES or endosomal recycling compartment (Farhan et al, 2017; Lamb et al, 2016; Sanchez-Wandelmer et al, 2015). In yeast, Sec16, Sec23 and Sec24 are required for autophagosome formation (Ishihara et al, 2001), while in mammalian cells ERES contributes to autophagy via a subset of COPII vesicles marked by particular isoforms of the inner shell proteins, SEC23B, SEC24A and SEC24B (Jeong et al, 2018).

Both degradative and biosynthetic membrane transport pathways undergo profound remodeling upon cell differentiation. Particularly rapid and intense membrane rearrangements occur during erythroid cell maturation. In human, erythroid progenitors undergo enormous expansion to fulfill the daily requirement of around 2 × 10^11^ new erythrocytes (Dzierzak & Philipsen, 2013). This robust differentiation involves efficient iron uptake through transferrin endocytosis, surface proteome remodeling, autophagic clearance of intracellular organelles, enucleation and elevated exocytosis (Moras et al, 2017; Ney, 2011).

A possible candidate protein that could be involved in rearrangement of membrane trafficking pathways during cell differentiation is bone morphogenetic protein 2 (BMP-2)-inducible kinase (BMP2K). It has been discovered as a transcriptional target of BMP-2 in osteoblast differentiation (Kearns et al, 2001) but later identified as an accessory protein of clathrin coated vesicles (Borner et al, 2012) and an interactor of an endocytic regulator NUMB (Krieger et al, 2013). Up to date, the cellular function of this member of the Ark1/Prk1 family of serine/threonine kinases is ill-defined and none of its phosphorylation targets are established. Its closest homologue, adaptor-associated kinase (AAK1) phosphorylates NUMB protein (Sorensen & Conner, 2008) and the medium (μ2) adaptin of the AP-2 complex (Conner & Schmid, 2002; Ricotta et al, 2002). Although proposed to regulate endosomal recycling (Henderson & Conner, 2007) and to enhance the clathrin assembly activity of AP-2 (Kadlecova et al, 2017), the role of AAK1 in cell biology is also not completely understood. While AAK1 may be important for neuronal cell function (Ultanir et al, 2012), very recent reports implicated BMP2K in the function of hematopoietic cells. The *BMP2K* gene was found as a novel fusion partner of ZNF384 oncogene and as a modulator of chemotherapy resistance in leukemia (Hirabayashi et al, 2017; Pandzic et al, 2016). It is also transcriptionally induced during *in vitro* stimulated erythropoiesis (Perucca et al, 2017). In a study that did not consider its possible endocytic functions, BMP2K was fished out as a putative stimulator of autophagy, potentially required for erythropoiesis (Potts et al. 2013). However, no insights were provided into possible mechanisms of how this putative endocytic kinase would regulate autophagy.

Here we report that splicing variants of BMP2K constitute a two-element system regulating transferrin endocytosis, SEC16A-dependent COPII trafficking and autophagy. Our study uncovers an unusual mechanism of two splicing variants of a kinase playing opposing roles in intracellular processes that may allow for their fine-tuning during cell differentiation.

## RESULTS

### The levels of Bmp2k splicing variants are initially increased but later reduced during mouse erythroid differentiation

To choose a proper biological context to study cellular functions of BMP2K, we mined gene expression databases and found that the human *BMP2K* gene is highly expressed in the early erythroid lineage (biogps.org). Moreover, using the Immunological Genome Project interface (immgen.org) we found that BMP2K mRNA levels are upregulated during human erythroid maturation in a manner similar to that of erythroid-enriched markers, such as TFRC (transferrin receptor 1) (Novershtern et al, 2011). To verify these data, we analyzed mRNA abundance of mouse Bmp2k in an *ex vivo* erythropoiesis model. According to the UniProtKB database, mouse expresses two splicing variants (isoforms) of the kinase, the longer (Bmp2k-L) and the shorter (Bmp2k-S), which result from alternative mRNA splicing. We observed that in isolated mouse fetal liver erythroblasts differentiated with erythropoietin (EPO)-containing medium, mRNA levels of Bmp2k-L and Bmp2k-S increased gradually, similarly to transferrin receptor 1 (Tfrc) (**Fig. S1A**). Consistently, similarly to Tfrc protein, the amounts of both variants were markedly elevated in differentiated cells (**Fig. 1A**).

**Figure 1.**
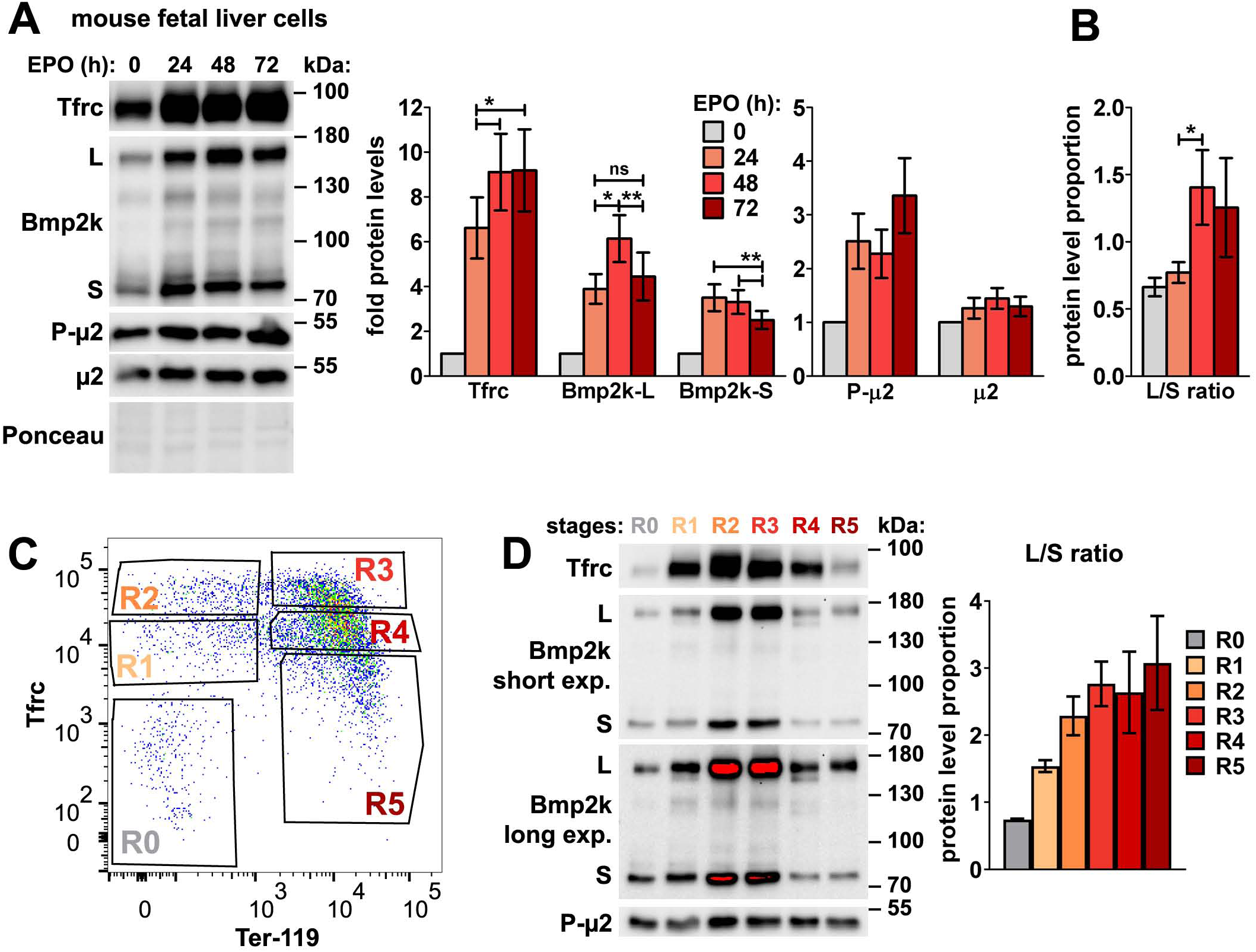
The levels of the longer (L) and the shorter (S) Bmp2k splicing variants as well as the L/S ratio are regulated during mouse erythroid differentiation. **A.** Representative western blots showing the levels of Tfrc, Bmp2k splicing variants (L and S) as well as total and Thr156-phosphorylated μ2 (P-μ2) at different time-points during erythropoietin (EPO)-stimulated differentiation of mouse fetal liver erythroblasts. Graphs show fold changes in non-normalized protein levels obtained by densitometric analysis of western blotting results (n=5 +/- SEM). **B.** The proportion between the detection intensities of BMP2K-L and -S (L/S ratio) calculated after densitometric analysis of bands from western blotting results represented in A (n=6 +/- SEM). **C.** Dot plot showing fluorescence intensities of the indicated markers found on the surfaces of mouse fetal liver erythroblasts stimulated with EPO for 96 h. Gates distinguish consecutive differentiation stages (R0-R5) of erythroblasts isolated by fluorescence activated cell sorting (FACS). **D.** Representative western blots showing the levels of Tfrc, Bmp2k variants as well as Thr156-phosphorylated μ2 (P-μ2) in the indicated FACS-isolated differentiation stages of erythroblasts. Graph shows the L/S ratio calculated after densitometric analysis of western blotting results (n=3 +/- SEM). *p<0.05, **p<0.01.

By analyzing consecutive time-points (24 h, 48 h and 72 h) of differentiation, we observed that after initial upregulation, the protein levels of both variants were subsequently downregulated (**Fig. 1A**). Noteworthy, the proportion between the intensities of western blotting detection of the two isoforms (L/S ratio) changed with time of differentiation, as Bmp2k-S protein was upregulated earlier (the highest levels detected at 24 h) than that of Bmp2k-L (the highest levels detected at 48 h) (**Fig. 1B**).

EPO-stimulated mouse fetal erythroblast cultures are a heterogeneous mixture of cells at various differentiation stages (Zhang et al, 2003). To assess precisely the amounts of Bmp2k variants at particular stages, we labelled EPO-stimulated cells with antibodies recognizing two commonly used mouse erythroid surface markers, CD71/Tfrc and Ter-119 (Zhang et al, 2003). This allowed us to isolate, by fluorescence activated cell sorting (FACS), the earliest primitive progenitors (CD71^low^/Ter-119^low^ – population R0) as well as further stages of erythroblast differentiation (consecutively: CD71^med^/Ter-119^low^ – R1, CD71^high^/Ter-119^low^ – R2, CD71^high^/Ter-119^high^ – R3, CD71^med^/Ter-119^high^ – R4 and CD71^low^/Ter-119^high^ – R5) (**Fig. 1C**). Consistently with the analysis of heterogeneous cultures (**Fig. 1B**), the levels of Bmp2k isoforms were low in the early stages (R0 and R1), the highest in the transitory stages (R2 and R3) and reduced at the last stages (R4 and R5) (**Fig. 1D**). Again, erythroid differentiation was associated with a change in the L/S ratio. It was in favor of Bmp2k-S (L/S < 1) in primitive progenitors (R0) but was increasingly shifted towards Bmp2k-L (L/S > 1) upon differentiation, even when total BMP2K protein levels were again downregulated (at R4 and R5) (**Fig. 1D**).

Thus, we found that during erythroid differentiation, the levels of BMP2K splicing variants are initially upregulated and subsequently reduced. The ratio between the two isoforms changes during erythroid differentiation, with Bmp2k-S being predominant in the early erythroid precursors and Bmp2k-L prevailing during differentiation and maturation.

### Reducing the levels of BMP2K splicing variants in K562 cells promotes erythroid differentiation accompanied by elevated transferrin endocytosis

The expression dynamics of mouse Bmp2k variants upon differentiation could suggest that the initial increase of their levels would promote early steps of erythropoiesis while their subsequent decrease would favor erythroid maturation. To verify this complex hypothetical scenario we sought a simpler cellular model. We found that K562 human erythroleukemia cells contain much higher amounts of BMP2K than various immortalized, solid tumor or non-erythroid blood cancer cell lines (**Fig. 2A and B**). In K562 cells the longer isoform (BMP2K-L) appeared as more abundant than the shorter (BMP2K-S) (**Fig. 2B, C and D**). Their L/S ratio was approximately 2:1 (calculated for control cells shown in **Fig. 2C and D**). This was reminiscent of R2 or R3 differentiation stages of mouse erythroblasts (**Fig. 1D**).

**Figure 2.**
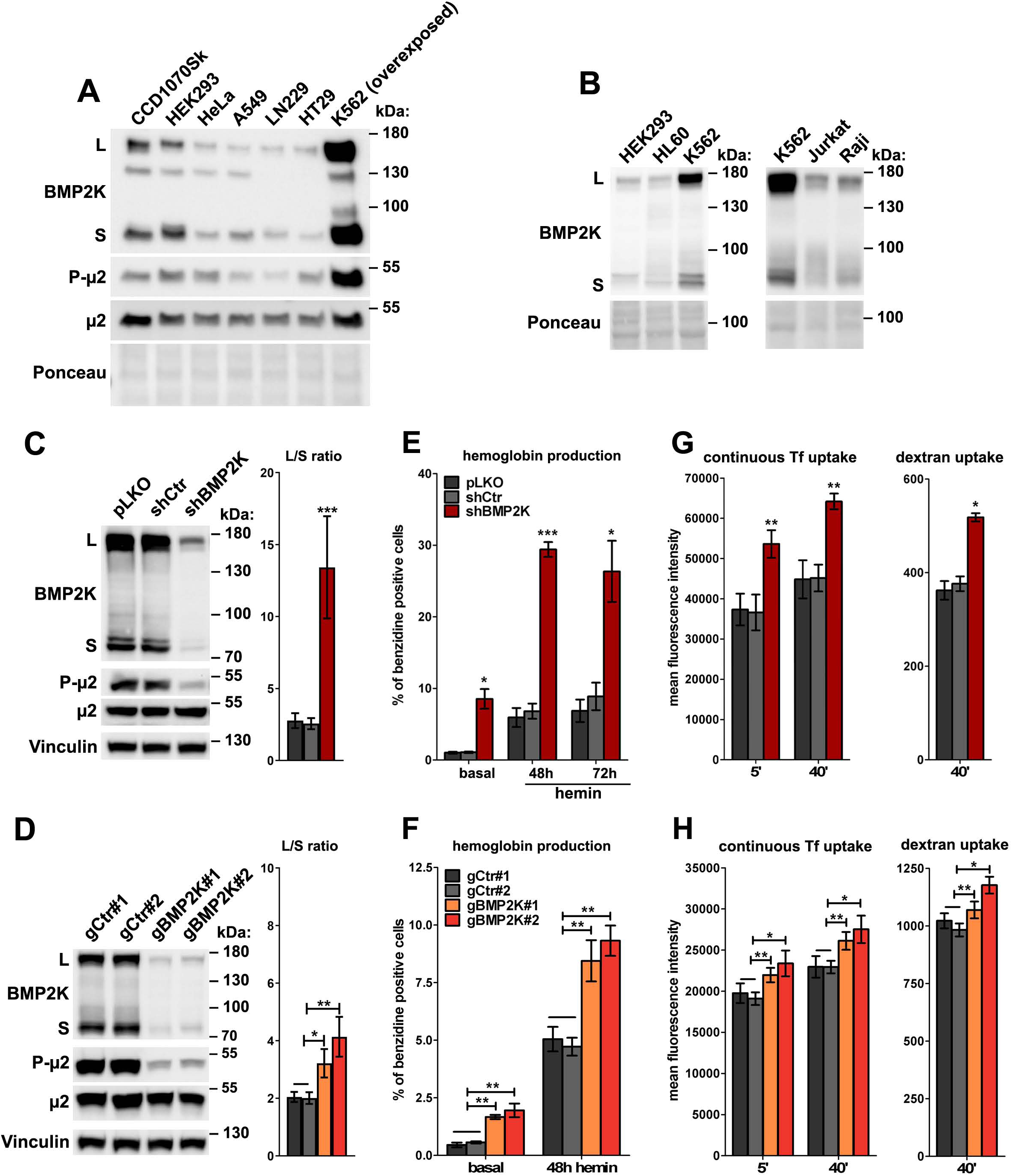
BMP2K splicing variants are enriched in K562 cells where they restrict erythroid differentiation and endocytic Tf uptake in a manner dependent on the L/S ratio. **A and B.** Western blots showing the levels of BMP2K splicing variants (L and S) or total and Thr156-phosphorylated μ2 (P-μ2) in lysates from the indicated human cell lines. Ponceau staining serves as a gel loading control. **C and D.** Western blots showing the efficiencies of depleting all BMP2K variants using shRNA (shBMP2K in C) or CRISPR/Cas9-mediated (gBMP2K#1 or #2 in D) approaches in K562 cells, and their effects on the levels of total and phosphorylated μ2, as compared to non-depleted cells. Control constructs used were: empty pLKO vector or non-targeting shRNA, shCtr (C) or non-targeting gRNAs, gCtr#1 and #2 (D). Graphs show the L/S ratio calculated after densitometric analysis of western blotting results (n=6 in C and 5 in D +/- SEM). **E and F.** Percentage of benzidine-positive control cells or cells depleted of BMP2K using shRNA (E) or CRISPR-Cas9 (F) approach, under basal growth conditions or after stimulation for 48 h or 72 h (A) or 72 h only (B) with hemin (20 μM). The quantification was performed using ImageJ software in 4-5 images of benzidine-stained cells (∼1000 cells) per analyzed condition (n=3 in E and 4 or 5 in F +/- SEM). **G and H.** Flow cytometry analysis of AlexaFluor647-Transferrin (Tf) or AlexaFluor488-dextran levels in control K562 cells or cells depleted of all BMP2K splicing variants using shRNA (G) or CRISPR/Cas9 (H) approach, after allowing for continuous Tf endocytosis at 37°C for 5’ or 40’, or after 40’ incubation with dextran at 37°C (n=3 or 4 in G and 4 in H +/- SEM). Percentage (E and F) as well as fluorescence intensity (G and H) values in BMP2K-depleted cells were compared statistically to those in shCtr-treated cells (E and G) or to (E) the average of values measured for gCtr#1 and gCtr#2-transduced cells (F and H). *p<0.05, **p<0.01, ***p<0.001.

K562 cells have erythroid progenitor-like features (Andersson et al, 1979) and can initiate erythroid differentiation, thus are widely used to study molecular aspects of erythropoiesis (Barbarani et al, 2017; Bu et al, 2014; Ma et al, 2013; Wang et al, 2011; Wu et al, 2018). To learn whether BMP2K silencing would reverse or advance erythroid differentiation of K562 cells, we silenced *BMP2K* gene expression using shRNA (shBMP2K) or CRISPR/Cas9 (gBMP2K#1 or #2) approaches. In each case, we achieved efficient reduction of BMP2K variant levels with a concomitant upregulation of the control 2:1 L/S ratio, very strongly (up to 13:1) upon shBMP2K and less potently (up to 3:1 or 4:1) upon gBMP2K#1 or #2 (**Fig. 2C and D**).

To assess the effects of BMP2K depletion on erythroid differentiation, we measured the expression levels of erythroid-specific genes, by qPCR, and the production of hemoglobin, by benzidine staining. As described (Villeval et al, 1983), control K562 cells expressed several erythroid-specific genes (**Fig. S1B and C**) but only 0.5%–1% of these cells were positive for hemoglobin (**Fig. 2E and F, S2A and B**). This percentage increased upon treatment with a heme precursor hemin, as described (Ma et al, 2013; Wang et al, 2011). We found that BMP2K depletion promoted erythroid differentiation of K562 cells, roughly proportionally to the observed L/S ratio. shBMP2K markedly elevated expression of erythroid-specific genes (**Fig. S1B**) and potently increased the number of hemoglobin-positive cells under basal culture conditions (by 6-fold) and in the presence of hemin (by 4-fold) (**Fig. 2E and S2A**). In turn, both gRNAs weakly induced the expression of erythroid markers (**Fig. S1C**) and stimulated hemoglobin production less potently than shBMP2K (∼3-fold in basal conditions and ∼2-fold upon hemin treatment) (**Fig. 2F and S2B**).

The elevated erythroid differentiation could result from increased transferrin (Tf) endocytosis (Moras et al, 2017). Thus, we measured by flow cytometry the effect of *BMP2K* silencing on continuous uptake of fluorescently labelled Tf and 10 kDa dextran (a fluid-phase marker) by K562 cells. shBMP2K increased early (5 min) and steady state (40 min) Tf uptake as well as dextran internalization, all by around 30% (**Fig. 2G, S3B**). gBMP2K#1 or #2 had a weaker effect on endocytosis than shBMP2K, increasing early and steady state Tf uptake roughly by 10%-15% and dextran internalization only by 5%-15% (**Fig. 2H, S3C**).

Hence, once BMP2K protein levels are high in erythroid cells, their reduction stimulates Tf endocytosis and favors red blood cell maturation, roughly proportionally to the resultant L/S ratio. This showed that a) at least one of the BMP2K variants inhibits cellular events responsible for erythroid maturation, and b) the balance between variant levels affects this inhibitory function.

### In K562 cells lacking all BMP2K variants, the elevated Tf uptake occurs despite reduced Thr156 µ2 phosphorylation, controlled by BMP2K-L

We reasoned that BMP2K variants could affect endocytosis via phosphorylation of µ2 adaptin on Thr156, suspected to be mediated by this kinase (Wrobel et al, 2019). Indeed, the elevated amounts of Bmp2k isoforms in mouse cells correlated with an increase in this phosphorylation (**Fig. 1A**). Moreover, K562 cells had high phospho-µ2 levels (**Fig. 2A**) that were strongly reduced upon BMP2K depletion (**Fig. 2C and D**). Although this phosphorylation was shown to promote endocytosis (Olusanya et al, 2001; Ricotta et al, 2002; Wrobel et al, 2019), the elevated continuous Tf uptake upon BMP2K depletion was associated with reduced phospho-µ2 levels (**Fig. 2G and H**).

To address whether one or both BMP2K variants regulate Tf endocytosis and µ2 phosphorylation, first we analyzed *in silico* the sequences of the human BMP2K-L and -S (**Fig. 3A**). They are identical at their N-termini, including a putative N-degron motif facilitating proteasomal degradation, and contain the serine-threonine protein kinase domain. BMP2K-L has 1161 amino acids, including 510 amino acids at its C-terminus absent in the shorter variant. BMP2K-S has 662 amino acids of which the first 650 are identical with BMP2K-L (**Fig. 3A**). Analyzing the sequences of the splicing variants in more detail, we found that both have EPS15 (NPF) and clathrin (DLL) binding motifs, while the L variant has one additional clathrin (SLL) and three AP-2 (DPF) binding motifs within its C-terminal tail (**Fig. 3A**). The presence of DPF motifs in BMP2K-L suggested that this variant could bind AP-2 and phosphorylate µ2. Consistently, using monoclonal antibodies recognizing all BMP2K splicing variants, we found that BMP2K-L, but not -S, co-precipitated with phosphorylated μ2 in K562 cells (**Fig. 3B**).

**Figure 3.**
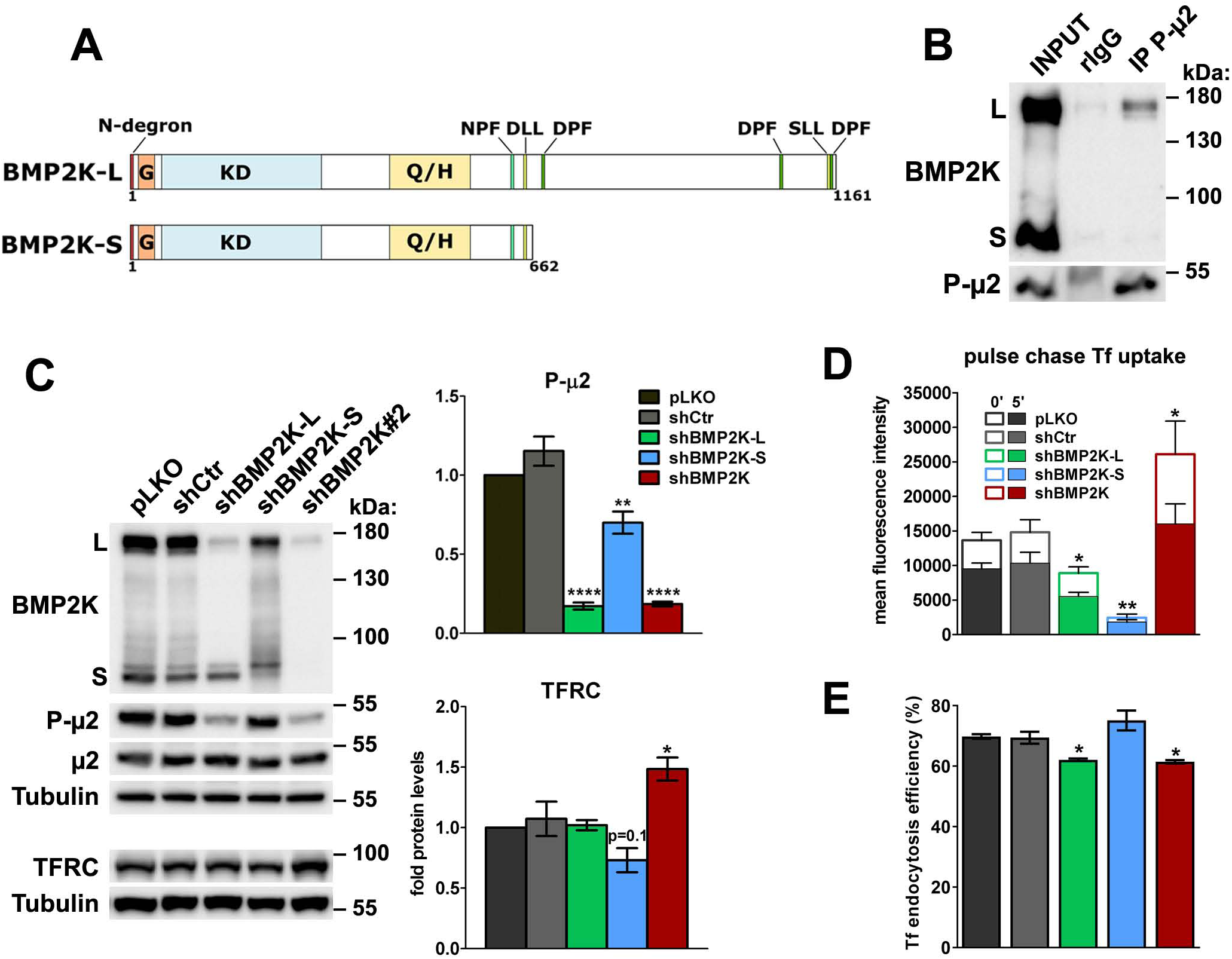
BMP2K-L promotes µ2 phosphorylation and CME efficiency, however cells lacking all BMP2K variants have an increased net Tf uptake. **A.** Schematic representation of BMP2K splicing variants containing glycine-rich (G), serine-threonine kinase (KD) and glutamine and histidine-rich (Q/H) domains as well as the indicated amino acid motifs. **B.** Western blots showing levels of endogenous BMP2K variants and Thr156-phosphorylated μ2 (P-μ2) in K562 cell lysates (INPUT) or samples after immunoprecipitation (IP) using either control immunoglobulins (rIgG) or antibodies recognizing phospho-μ2. **C.** Western blots showing shRNA-mediated depletion of single (shBMP2K-L or shBMP2K-S) or of all BMP2K variants (shBMP2K) and their impacts on Thr156 phosphorylation of μ2 and on the levels of μ2 and TFRC proteins, as compared to empty pLKO vector or non-targeting shRNA construct, shCtr. Graphs show densitometric analysis of western blotting bands for the indicated proteins using tubulin abundance for normalization (n=4 +/- SEM). **D.** Pulse-chase Tf uptake in cells with depletion of BMP2K variants. The empty bars represent the amount of fluorescently-labelled Tf pre-bound to cells and the solid bars show the amount of Tf internalized at 37°C for 5’ (n=3 +/- SEM). **E.** The percentage of internalized Tf with respect to the amount of pre-bound Tf (n=3 +/- SEM).

Next, we designed shRNAs to reduce the expression specifically of BMP2K-L or BMP2K-S. Silencing of the longer variant had no effect on the expression of the shorter, however BMP2K-S depletion modestly reduced full length BMP2K-L levels (**Fig. 3C**). As expected, μ2 phosphorylation was strongly reduced upon knock-down of the L isoform and only modestly downregulated by silencing of the S variant (**Fig. 3C**). In cells depleted of both isoforms (shBMP2K), the remaining cellular levels of the longer variant as well as reduction of μ2 phosphorylation were the same as in shBMP2K-L-treated cells (**Fig. 3C**).

Thus, we found that in K562 cells lacking all *BMP2K* gene products, the elevated Tf uptake occurs despite reduced Thr156 phosphorylation of µ2 which is under control of BMP2K-L but not BMP2K-S. This suggested that BMP2K variants might be differentially implicated in AP-2-mediated CME.

### The role of BMP2K splicing variants in CME does not explain their involvement in erythroid differentiation

To investigate the role of BMP2K variants in CME, we applied a pulse-chase endocytosis assay (**Fig. 3D and E, S3D**). Therein the amount of fluorescently-labelled Tf pre-bound to cells on ice (0’, empty bars in **Fig. 3D**) prior to endocytosis reflects cell surface abundance of Tf receptors, while the percentage of Tf internalized at 37°C (5’, solid bars in **Fig. 3D**) with respect to the pre-bound Tf represents CME efficiency (**Fig. 3E**). BMP2K-L depletion and to an even higher extent BMP2K-S depletion reduced the amount of Tf bound to the cell surface (by 35% and by 80%, respectively) (**Fig. 3D**). This was not due to altered total TFRC protein levels as they were unchanged upon shBMP2K-L and only modestly reduced upon shBMP2K-S (**Fig. 3C**). Hence, the BMP2K variants could in part affect Tf uptake through the regulation of Tf receptor availability on the cell surface that had to be taken into account for the proper interpretation of the pulse-chase endocytosis assay.

In control K562 cells, 5’ pulse-chase uptake resulted in internalization of 70% of the pre-bound Tf (**Fig. 3D**). As expected based on the presence of DPF motifs in BMP2K-L, depletion of this variant reduced the percentage of internalized Tf down to 62%, hence moderately inhibiting CME efficiency (**Fig. 3E**). Overall, due to the weaker Tf surface binding and the lower CME efficiency, BMP2K-L-depleted cells internalized almost twice less Tf than control cells (**Fig. 3D**). BMP2K-S-depleted cells, despite showing a tendency for an increased endocytosis rate (**Fig. 3E**), overall internalized very little Tf (**Fig. 3D**).

The analysis of CME efficiency in cells depleted of single BMP2K isoforms did not provide explanation for the elevated Tf uptake in cells lacking all *BMP2K* gene products due to shBMP2K (shown in **Fig. 2G)**. These cells had elevated TFRC mRNA and total protein levels (**Fig. S1B and 3C**) and showed a two-fold increase in Tf surface binding (**Fig. 3D**), arguing for a higher number of Tf receptor molecules available on the PM. Hence, although they showed the same reduction of Tf internalization efficiency (down to 62%) as shBMP2K-L-treated cells (**Fig. 3E**), their resultant Tf pulse-chase uptake was higher than in control cells (**Fig. 3D**). This indicated that the elevated Tf uptake upon depletion of all BMP2K isoforms by shBMP2K (**Fig. 2G)** occurs at least in part due to higher abundance of Tf receptor molecules on the cell surface, despite the lower Tf CME efficiency.

Collectively, although we confirmed the implication of BMP2K kinase in endocytosis, and specifically a stimulatory role of BMP2K-L, the functions of BMP2K variants in Tf internalization do not correlate with their effects on differentiation. This suggested that in addition to endocytosis, BMP2K variants must regulate other cellular events to affect red blood cell maturation.

### BMP2K-S, in addition to CME adaptors, associates with regulators of the early secretory pathway

To find out in which other cellular processes the two BMP2K isoforms could function, we investigated their interactomes. To this end, we performed proximity biotinylation (BioID) followed by liquid chromatography coupled to tandem mass spectrometry (LC-MS/MS) in HEK293 cells ectopically expressing the L or the S variant tagged with a mutant BirA biotin ligase (BirA*) at their N- or C-termini (**Fig. S4A and B**). Most likely due to the presence of the N-degron motif, the C-terminally tagged proteins were expressed less efficiently than those tagged N-terminally (**Fig. S4B**). To avoid artifacts due to either very high expression of N-terminally tagged proteins or N-degron-mediated degradation of C-terminally tagged variants, we focused on proteins detected as common interactors of N- and C-terminally tagged BMP2K isoforms (**Table S1 and S2, Fig. 4A, S4C**). Within the BMP2K-L interactome, we found proteins involved in CME as well as mRNA translation and cotranslational protein targeting to membranes, while BMP2K-S was found to interact with proteins acting in CME, ER-Golgi transport, RNA export through nuclear pores and mRNA translation, decay and transport (**Fig. S4C**).

**Figure 4.**
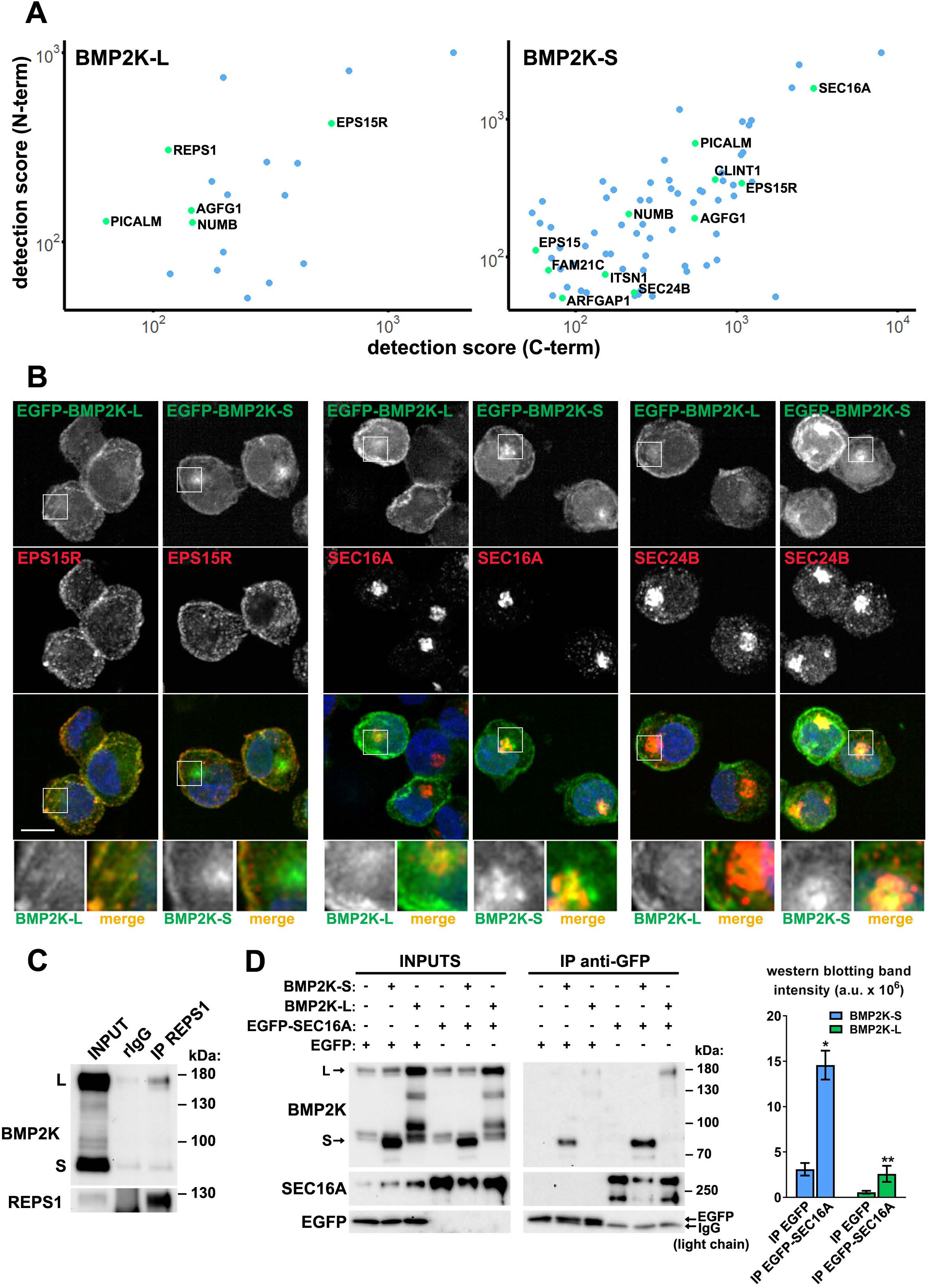
The BioID interactome analysis and its subsequent validation show that both BMP2K splicing variants can interact with SEC16A. **A.** Dot plots showing BioID-MS detection scores (log scale) of proteins found as proximal to both N- and C-terminally tagged BMP2K-S or -L variants in HEK293 cells. **B.** Maximum intensity projection images from confocal microscope showing localization of ectopically expressed EGFP-tagged BMP2K-L or -S with respect to the indicated proteins and cell nuclei marked with DAPI stain (blue) in K562 cells. Insets: Magnified views of boxed regions in the main images. Scale bars, 10 µm. **C.** Western blots showing levels of endogenous BMP2K and REPS1 in K562 cell lysates (INPUT) or samples after immunoprecipitation (IP) using either control immunoglobulins (rIgG) or antibodies recognizing REPS1 protein. **D.** Levels of BMP2K-L, BMP2K-S, EGFP or EGFP-tagged SEC16A in whole cell lysates (INPUTS) or in immunoprecipitates (IP) using anti-GFP antibodies from HEK293 cells. Different combinations of simultaneous ectopic expression of the analyzed proteins are indicated above the images. Graph shows non-normalized densitometric analysis of western blotting bands expressed in arbitrary units (a.u.; n=3 +/- SEM).

In the BioID analysis, we did not detect clathrin proteins and AP-2 complex components. However, among endocytic regulators found as proximal to both BMP2K isoforms were established adaptors of clathrin coated pits and/or AP-2 interactors: NUMB, PICALM, EPS15R, and AGFG1 (HRB) (Benmerah et al, 1995; Chaineau et al, 2008; Coda et al, 1998; Miller et al, 2015; Santolini et al, 2000; Tebar et al, 1996) (**Fig. 4A and S4C**). Reassuringly, NUMB has been previously experimentally confirmed as an interactor of BMP2K (Krieger et al, 2013). The only vesicular trafficking-related protein identified by our BioID as specific to BMP2K-L was REPS1 that, likewise BMP2K, has been described as a component of clathrin-coated vesicles (Borner et al, 2012) and a NUMB interactor (Krieger et al, 2013). In turn, most of trafficking-related proteins identified as proximal to BMP2K-S and not detected for the L isoform (i.e. SEC16A, SEC24B, ARFGAP1, EPS15 and CLINT1) could be annotated either to ER-Golgi transport or trafficking from the Golgi to the endolysosomal system (Chi et al, 2008; Mills et al, 2003; Watson et al, 2006; Wendeler et al, 2007; Yang et al, 2002) (**Fig. 4A and S4C**). Of note, SEC16A was detected with the highest MS score among all trafficking regulators identified for BMP2K-S (**Fig. 4A).** Collectively, the BioID analysis indicated that BMP2K splicing variants might differentially associate with REPS1 (BMP2K-L) or markers of ER-Golgi trafficking (BMP2K-S).

### Both BMP2K splicing variants can associate with SEC16A protein

To verify the BioID results, we first asked whether BMP2K isoforms localized intracellularly to endocytic and/or secretory compartments. In K562 cells, using the monoclonal antibody recognizing in western blotting all BMP2K variants, by confocal microscopy we detected signal predominantly near the PM. There, it overlapped to some extent with EPS15R, found in BioID as proximal to both isoforms (**Fig. S4D**). However, intensity of the signal detected with anti-BMP2K antibody, although weaker due to BMP2K-L depletion, did not decline upon shBMP2K-S (**Fig. S4D and E**). Moreover, in cells depleted of all BMP2K variants, the signal reduction, as compared to control cells, was similar to cells lacking BMP2K-L only (**Fig. S4D and E**). Thus, considering that the monoclonal antibody was not suitable for detection of BMP2K-S in microscopy, we analyzed the intracellular localization of EGFP-tagged BMP2K variants ectopically expressed in K562 cells. As expected, EGFP-BMP2K-L was enriched predominantly near the PM, where it colocalized with EPS15R (**Fig. 4B and S5A**). BMP2K-S, although also overlapping with EPS15R near the PM, strongly concentrated in the juxtanuclear region, positive for SEC16A or SEC24B (**Fig. 4B and S5A**). Such juxtanuclear localization to SEC16A or SEC24B-positive compartment, however much weaker, could also be observed for EGFP-BMP2K-L (**Fig. 4B and S5A**).

The BioID analysis suggested that REPS1 is a specific interactor of BMP2K-L. Consistently, in K562 cells endogenous BMP2K-L, but not -S, co-precipitated with REPS1 (**Fig. 4C**). Conversely, SEC24B, identified by BioID as a specific BMP2K-S interactor, despite very efficient immunoprecipitation did not co-precipitate any of the endogenous BMP2K variants in K562 cells (**Fig. S5B**). Unfortunately, we were not able to efficiently immunoprecipitate endogenous SEC16A protein likely due to its large size and lack of appropriate antibodies. Hence, to assess the ability of BMP2K-L or -S to associate with SEC16A, we overexpressed them in HEK293 cells together with EGFP only or with EGFP-tagged SEC16A, and performed immunoprecipitation using anti-GFP antibodies. Although we repeatedly observed some non-specific co-precipitation of both BMP2K isoforms on agarose resin with EGFP only, they were significantly enriched in EGFP-SEC16A precipitates (**Fig. 4D**).

Collectively, we confirmed the BioID findings that REPS1 is a BMP2K-L-specific interactor and that SEC16A can interact with both BMP2K splicing variants.

### In K562 cells, SEC16A differentially regulates production of juxtanuclear and dispersed COPII vesicles

We assumed that SEC16A would be key to delineate the non-endocytic functions of BMP2K variants. SEC16A regulates production of COPII vesicles (Glick, 2014; Lord et al, 2013), whose trafficking contributes to protein secretion or autophagic degradation (Farhan et al, 2017), both important for erythropoiesis (Grosso et al, 2017; Satchwell et al, 2013). However, the role of SEC16A in erythroid cells has not been studied. Thus, we first needed to verify SEC16A function in COPII trafficking and autophagy in K562 cells, before elucidating its interplay with BMP2K variants.

To test SEC16A role in COPII trafficking, we analyzed its intracellular distribution together with SEC31A, a ubiquitously expressed mammalian Sec31 homologue (D’Arcangelo et al, 2013; Satchwell et al, 2013; Tang et al, 2000). As expected from studies in other cell types (Martinez-Menarguez et al, 1999; Orci et al, 1991; Stephens et al, 2000), both SEC16A and SEC31A-positive structures concentrated in the juxtanuclear region but were also dispersed throughout the cytoplasm (pLKO and shCtr control cells in **Fig. 5A**). The juxtanuclear SEC16A compartment was juxtaposed to ERGIC and cis-Golgi, marked by a component of COPI assemblies ARFGAP1 (Yang et al, 2002) (**Fig. S5C**), that in turn were in close proximity to juxtanuclearly located TGN, identified by TGN46 immunostaining (**Fig. S5D**). Hence, given the text-book compartmentalization of early secretory organelles, the juxtanuclear SEC16A compartment constituted ERES in K562 cells (Lord et al, 2013). The staining intensity of SEC31A-positive vesicular structures (mean fluorescence intensity per vesicle in **Fig. 5B**, hereafter referred to as SEC31A load) was higher in the juxtanuclear than in the dispersed COPII vesicular structures. Based on intracellular localization and SEC31A load we considered the juxtanuclear structures as functional secretory COPII vesicles.

**Figure 5.**
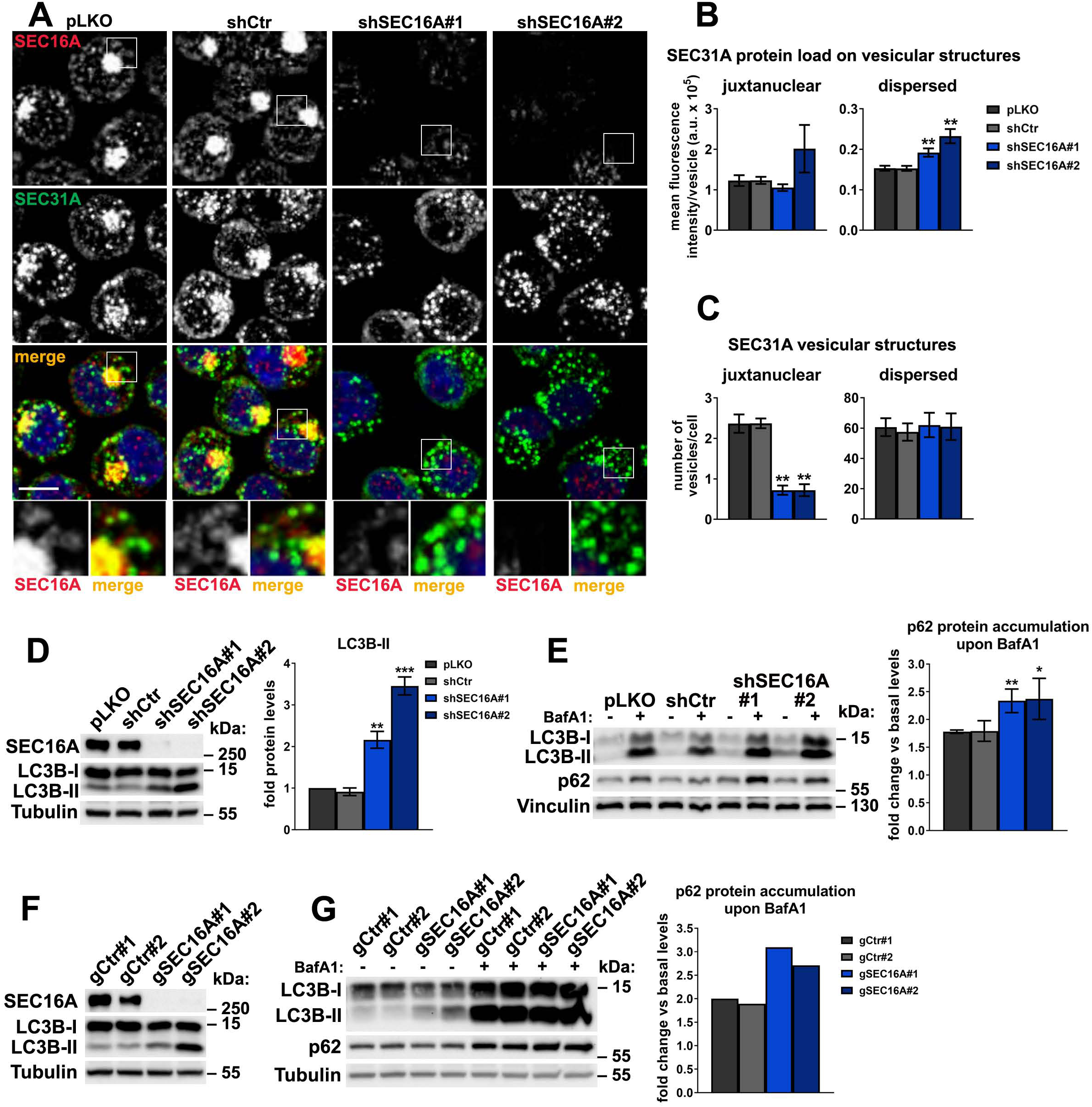
In K562 cells, SEC16A differentially regulates generation of juxtanuclear and dispersed COPII vesicles and limits autophagic degradation. **A.** Representative maximum intensity projection images from confocal microscope, showing immunolocalization of SEC16A and SEC31A proteins as well as cell nuclei marked with DAPI stain (blue) in control (pLKO, shCtr) or SEC16A-depleted (shSEC16A#1 and #2) K562 cells. Insets: Magnified views of boxed regions in the main images. Scale bar, 10 µm. **B and C.** SEC31A mean fluorescence intensity per vesicle (SEC31A protein load) within juxtanuclear or dispersed structures, presented in arbitrary units (a.u. in B) and the number of juxtanuclear or dispersed SEC31A-positive vesicles per cell (C) in control or SEC16A-depleted cells. Quantification from images represented by those in A (n=4 +/- SEM). **D and F.** Western blots showing the efficiencies of shRNA (D) or CRISPR/Cas9-mediated (F) SEC16A depletion in K562 cells and their effects on the levels of LC3B-I, LC3B-II and p62 proteins in K562 cells. Graph in D shows densitometric analysis of western blotting bands for the indicated proteins using tubulin abundance for normalization (n=3 +/- SEM). **E and G.** The abundance of p62 protein in non-treated cells or in cells treated for 15 h with 75 nM bafilomycin A (BafA1). Graphs show calculated fold increase of p62 protein levels caused by BafA1, obtained after densitometric analysis of western blotting bands using vinculin abundance for normalization (n=4 +/- SEM in E and n=1 in G). Fold protein levels in cells depleted of SEC16A (D, E) were compared statistically to the levels in cells treated with shCtr. *p<0.05, **p<0.01, ***p<0.001.

To verify whether in K562 cells SEC16A is required for COPII trafficking, we depleted it using two individual shRNAs (shSEC16A#1 and #2) (**Fig. 5A**). Both led to a loss of juxtanuclear COPII vesicles (strongly reduced number of SEC31A vesicles, **Fig. 5C**), consistent with requirement of SEC16A for COPII assembly (Martinez-Menarguez et al, 1999; Watson et al, 2006). However, the absence of SEC16A protein did not affect the number of the dispersed SEC31A structures (**Fig. 5C**) but increased their SEC31A protein load (**Fig. 5B**). This indicated that in the dispersed COPII structures, SEC16A could inhibit SEC31A recruitment, reminiscent of the function described for yeast Sec16 (Bharucha et al, 2013; Kung et al, 2012; Yorimitsu & Sato, 2012).

Hence, we found that in K562 cells SEC16A is indispensable for the generation of juxtanuclear secretory COPII vesicles but not dispersed COPII assemblies, where it may negatively regulate SEC31A recruitment.

### In K562 cells, SEC16A limits basal autophagic flux

Having confirmed the role of SEC16A in regulation of COPII trafficking, we addressed whether it also regulates autophagic degradation in K562 cells. Although yeast Sec16 is required for autophagosome formation (Ishihara et al, 2001), to date such requirement has not been proven for mammalian SEC16A. To address this, we used cells lacking SEC16A to analyze the processing of LC3B-I to LC3B-II (LC3B lipidation), one of the indicators of the autophagy status (Mizushima et al, 2010). Surprisingly, shRNA-mediated knock-down of SEC16A elevated the LC3B lipidation (**Fig. 5D**), suggesting activated autophagy. To confirm this, we performed autophagic flux analysis (**Fig. 5E**). We inhibited autophagic degradation using bafilomycin A1 (BafA1) and assessed the accumulation of non-degraded SQSTM1/p62 protein (hereafter referred to as p62), an established cargo for autophagic degradation. In cells lacking SEC16A, the accumulation of p62 protein was higher than in control cells, demonstrating an elevated autophagic flux (**Fig. 5E**). As this finding was not accordant with findings for yeast Sec16, we verified it by analyzing the autophagy status of cells depleted of SEC16A by CRISPR/Cas9. Again, when compared to control cells, two single gRNAs targeting *SEC16A* gene (gSEC16A#1 and gSEC16A#2) increased LC3B lipidation (**Fig. 5F**) and the autophagic flux (**Fig. 5G**).

Collectively, we discovered that SEC16A not only regulates COPII trafficking but also limits basal autophagic degradation in K562 cells. This provided a basis to study the interplay between BMP2K splicing variants and SEC16A functions.

### BMP2K-L promotes while BMP2K-S limits production of COPII vesicles

To address whether BMP2K splicing variants could regulate SEC16A-dependent COPII trafficking or autophagy, we first analyzed whether shRNA-mediated depletion of BMP2K-L or -S had any effect on SEC16A abundance or distribution in K562 cells. shBMP2K-L led to a modest but reproducible increase, while shBMP2K-S reduced SEC16A protein levels (**Fig. 6A**). Depletion of all BMP2K isoforms (shBMP2K) lowered SEC16A levels, to the same extent as shBMP2K-S (**Fig. 6A**). By comparison, protein levels of SEC24B, another ER-Golgi transport-related protein identified in our BioID analysis, were essentially non-affected by depletion of BMP2K variants (**Fig. 6A**). To ensure that the observed changes in SEC16A abundance were not caused by RNAi off-target effects, we tested additional shRNAs. Reassuringly, shBMP2K-L#2 increased while shBMP2K-S#2 reduced SEC16A amounts (**Fig. S6A**). The regulation of SEC16A protein levels by the BMP2K-L or -S variants was not due to altered gene transcription. We only observed ∼30% downregulation of SEC16A mRNA levels in cells lacking all BMP2K variants (**Fig. 6B**).

**Figure 6.**
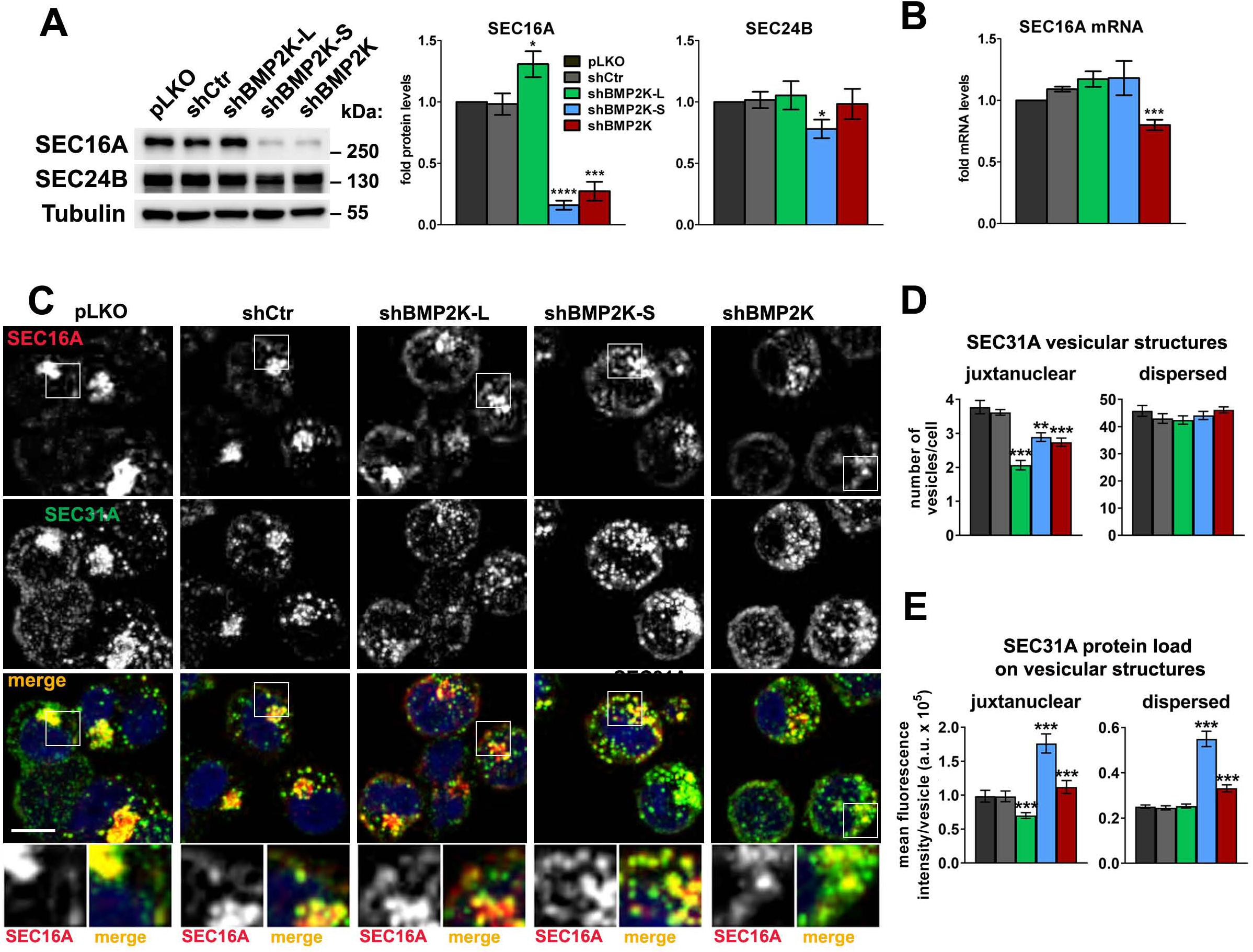
BMP2K-L promotes while BMP2K-S limits production and SEC31A load of COPII vesicles. **A.** The effect of shRNA-mediated depletion of single (shBMP2K-L or shBMP2K-S) or all BMP2K splicing variants (shBMP2K) on the levels of SEC16A and SEC24B proteins, as compared to empty pLKO vector or non-targeting shRNA construct, shCtr. Graphs show densitometric analysis of western blotting bands for the indicated proteins using tubulin abundance for normalization (n=5 +/- SEM). **B.** SEC16A mRNA fold levels in control cells or in cells with shRNA-mediated depletion of BMP2K variants (n=5 +/- SEM). **C.** Representative maximum intensity projection images from confocal microscope, showing the effect of shRNA-mediated depletion of BMP2K variants on immunolocalization of SEC16A and SEC31A proteins in K562 cells. Cell nuclei marked with DAPI stain (blue). Insets: Magnified views of boxed regions in the main images. Scale bar, 10 µm. **D and E.** The number of juxtanuclear and dispersed SEC31A-positive vesicles per cell (D) or SEC31A mean fluorescence intensity per vesicle (SEC31A protein load) presented in arbitrary units (a.u. in E) in control cells or cells lacking BMP2K variants. Quantification from images represented by those in C (n=5 +/- SEM). *p<0.05, **p<0.01, ***p<0.001, ****p<0.0001.

Depletion of BMP2K-L had no apparent effect on the morphology or staining intensity of juxtanuclear SEC16A structures but increased integral fluorescence intensity of the dispersed structures, arguing for their expansion (**Fig. 6C and S6B**). shBMP2K-S led to a diffusion of SEC16A-positive compartment, with lower staining intensity of juxtanuclear structures (**Fig. 6C and S6B**). Similarly, in cells lacking all BMP2K variants, the juxtanuclear SEC16A-compartment was also visually diffused but the SEC16A staining intensity was overall lower (**Fig. 6C and S6B**).

The bidirectional regulation of SEC16A protein levels and different effects on SEC16A distribution suggested that the two BMP2K variants could affect ERES function in an opposite manner. To verify this, we analyzed the effects of their depletion on distribution of SEC31A vesicles (**Fig. 6C**). Upon shBMP2K-L, the number (**Fig. 6D)** and SEC31A load (**Fig. 6E)** of juxtanuclear COPII vesicles decreased while those of dispersed structures did not increase despite the described above (**Fig. S6B**) expansion of dispersed SEC16A compartment. Consistently, total colocalization between SEC31A and SEC16A was lower than in control cells (**Fig. S6C).** This indicated less efficient COPII production and SEC31A recruitment at SEC16A-positive ERES in the absence of BMP2K-L. Conversely, shBMP2K-S, although had no clear effect on the number of COPII vesicles (**Fig. 6D)**, led to a strong increase in SEC31A protein load in both, juxtanuclear and dispersed structures (**Fig. 6E)** and higher total SEC31A/SEC16A colocalization (**Fig. S6C**). This pointed to a stimulated SEC31A recruitment at SEC16A-positive ERES.

Similarly to shBMP2K-S, shBMP2K had a negligible effect on the number of COPII vesicles (**Fig. 6D)**. Importantly, it significantly elevated SEC31A protein recruitment to COPII vesicles but to a lesser extent than upon shBMP2K-S (**Fig. 6E**). Of note, shBMP2K-treated cells showed reduced total SEC31A/SEC16A colocalization due to overall lower SEC16A staining intensity (**Fig. S6C**). As shown in **Fig. 3C**, shBMP2K-treated cells have the shorter isoform depleted to the same levels as upon shBMP2K-S but additionally contain lower amounts of the longer. Thus, by comparing the effects of shBMP2K-S and shBMP2K on SEC31A distribution, we concluded that the elevated recruitment of SEC31A into COPII structures in the absence of BMP2K-S is roughly proportional to the remaining BMP2K-L levels.

Collectively, we discovered that, likely through differential effects on SEC16A function, BMP2K-L promotes while BMP2K-S limits COPII vesicle production, including SEC31A recruitment. Hence, we identified a novel intracellular regulatory system, termed the BMP2K-L/S system, where the two splicing variants act in an opposite manner in controlling COPII trafficking.

### The BMP2K-L/S system controls the trafficking of SEC24B-positive COPII vesicles

To further verify the role of the BMP2K-L/S system, we analyzed the intracellular distribution of another COPII component, SEC24B upon depletion of BMP2K variants (**Fig. 7A**). We found SEC24B as potentially proximal to BMP2K-S (**Fig. 4A and S4C**), while yeast Sec24 was shown to modulate the inhibitory role of Sec16 on Sec31 recruitment (Kung et al, 2012).

**Figure 7.**
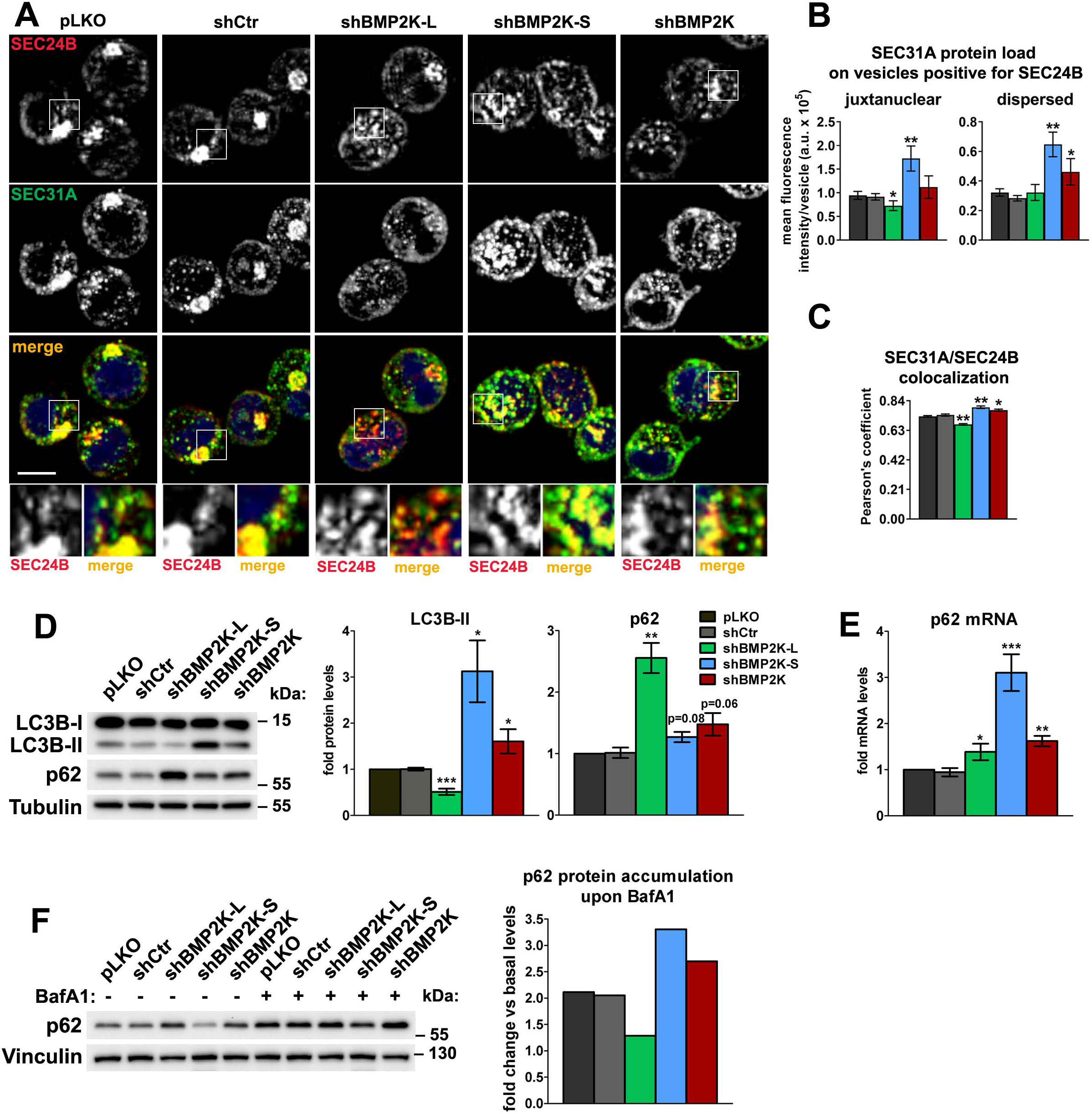
The BMP2K-L/S system controls the trafficking of SEC24B-positive COPII vesicles and autophagic flux. **A.** Representative maximum intensity projection images from confocal microscope, showing the effect of shRNA-mediated depletion of single (shBMP2K-L or shBMP2K-S) or all BMP2K splicing variants (shBMP2K) on immunolocalization of SEC24B with respect to immunostained SEC31A protein in K562 cells. Cell nuclei marked with DAPI stain (blue). Insets: Magnified views of boxed regions in the main images. Scale bar, 10 µm. **B and C.** SEC31A mean fluorescence intensity (SEC31A protein load) of vesicles positive for SEC24B, presented in arbitrary units (a.u. in B) and total colocalization between SEC31A and SEC24B structures quantified using the Pearson’s correlation coefficient (C) in control cells or cells lacking BMP2K variants. Quantification based on images as those represented in A (n=4 +/- SEM). **D.** Western blots showing the effects of depletion of BMP2K variants on the levels of LC3B-I, LC3B-II and p62 proteins in K562 cells. Graphs show densitometric analysis of western blotting bands for the indicated proteins using tubulin abundance for normalization (n=4 +/- SEM). **E.** p62 mRNA fold change levels in control cells or in cells with shRNA-mediated depletion of BMP2K variants (n=5 +/- SEM). **F.** The abundance of p62 protein in non-treated cells or in cells treated for 15 h with 75 nM bafilomycin A (BafA1). Graph shows calculated fold increase of p62 protein levels induced by BafA1, obtained after densitometric analysis of western blotting bands using vinculin abundance for normalization (n=1). *p<0.05, **p<0.01, ***p<0.001.

Depletion of BMP2K-L reduced integral intensity of SEC24B staining in the juxtanuclear but not in the dispersed structures (**Fig. S7**), consistently with the lower amount of juxtanuclear but not dispersed COPII vesicles (**Fig. 6D**). Moreover, cells lacking BMP2K-L had a reduced SEC31A load in juxtanuclear SEC24B-positive structures (**Fig. 7B**) and a lower total SEC31A/SEC24B colocalization (**Fig. 7C**). Thus, we found that BMP2K-L promotes production of and SEC31A recruitment onto juxtanuclear SEC24B vesicles. BMP2K-S depletion led to a diffusion of SEC24B compartment (**Fig. S7**) as it was the case for SEC16A (**Fig. S6C**). Consistently with the described above (**Fig. 6E**) strongly elevated SEC31A recruitment to ERES, cells lacking BMP2K-S showed increased SEC31A load on SEC24B-positive vesicles (**Fig. 7B**) and therefore higher total SEC31A/SEC24B colocalization (**Fig. 7C**). As shown in **Fig. 6D** and **E**, the BMP2K-L/S system regulates incorporation of SEC31A into COPII in a manner roughly proportional to the amount of the longer isoform. Consistently, shBMP2K increased SEC31A load in SEC24B-positive vesicles (**Fig. 7B**) and total SEC31A/SEC24B colocalization (**Fig. 7C**), but to a lower extent than shBMP2K-S.

Hence, we discovered that BMP2K variants regulate generation of COPII vesicles containing SEC24B. This reinforced our notion that the BMP2K-L/S system controls COPII trafficking and suggested that this system could regulate other COPII-dependent processes, such as autophagy.

### BMP2K-L promotes while BMP2K-S restricts autophagic degradation

Upon SEC16A depletion, higher recruitment of SEC31A into dispersed vesicular structures coincided with elevated autophagic flux (**Fig. 5**). Moreover, the BMP2K-L/S system regulated SEC31A recruitment to both, juxtanuclear and dispersed COPII vesicles (**Fig. 6C, D and E**), including those containing SEC24B (**Fig. 7A, B and C**), one of COPII components shown to promote autophagy (Jeong et al, 2018).

Although BMP2K was fished out as a stimulator of LC3-dependent autophagy (Potts et al, 2013), our observations of differential effects of BMP2K-L or -S on COPII production suggested that BMP2K variants could affect autophagy in an opposite manner. Consistently, shBMP2K-L reduced, while shBMP2K-S increased, the levels of lipidated LC3B (**Fig. 7D**), indicative of inhibited or activated autophagy, respectively. shBMP2K upregulated the abundance of LC3B-II, but to a lesser extent than BMP2K-S depletion (**Fig. 7D**). This intermediate effect of global BMP2K silencing, pointing to the induction of autophagy, could result from opposing actions of the two isoforms.

To validate the above results we analyzed autophagic degradation of p62. We observed that cells lacking BMP2K-L, although having unchanged p62 mRNA levels (**Fig. 7E**), showed a strong increase of its protein content (**Fig. 7D)**, likely due to restrained autophagy. Conversely, cells lacking BMP2K-S, despite a strong increase in p62 mRNA levels (**Fig. 7E**), did not accumulate the protein (**Fig. 7D**), which could be explained by elevated autophagic degradation. These observations reinforced the opposing effects of both BMP2K variants on autophagy, demonstrated by the analysis of LC3B lipidation. In turn, cells treated with shBMP2K showed slightly elevated levels of both p62 protein and mRNA (**Fig. 7D and E**), a phenotype difficult to interpret unequivocally.

To clarify the effect of depleting all BMP2K splicing variants on autophagy, we performed autophagic flux analysis. To this end, we inhibited autophagic degradation using BafA1 and analyzed the accumulation of non-degraded p62 protein in BMP2K-depleted cells (**Fig. 7F**). Consistent with autophagy inhibition, in cells lacking BMP2K-L, the accumulation of p62 protein was weaker than in control cells. In turn, in BMP2K-S-depleted cells, stronger p62 accumulation confirmed the increased autophagic flux. Importantly, silencing of all BMP2K splicing variants using shBMP2K also elevated p62 accumulation upon BafA1 treatment, indicative of increased autophagic flux (**Fig. 7F**). Thus, the net outcome of using shBMP2K, which efficiently removes the BMP2K-S and leaves a certain amount of BMP2K-L, is elevated basal autophagic degradation.

We thus discovered that the two BMP2K isoforms play opposite roles in regulation of autophagy. Collectively, we found that BMP2K-L, which promotes COPII trafficking, stimulates autophagy, while BMP2K-S, which inhibits COPII trafficking, limits autophagic degradation.

## DISCUSSION

A growing body of evidence suggests that AAK1 and BMP2K serine-threonine kinases might be of high importance for human physiology, with AAK1 implicated in functioning of the central nervous system (Kostich et al, 2016; Ultanir et al, 2012) and BMP2K involved in leukemogenesis (Hirabayashi et al, 2017; Pandzic et al, 2016; Wang et al, 2020). Although AAK1 is an established regulator of CME (Agajanian et al, 2019; Conner & Schmid, 2002; Loi et al, 2016; Ricotta et al, 2002; Sorensen & Conner, 2008), the cellular functions of BMP2K are largely unknown. Here we discover that its two erythroid-enriched splicing variants play different roles in processes crucial for erythroid differentiation such as endocytosis, COPII-dependent trafficking and autophagy (**Fig. 8**). Given a high amino acid homology (Sorrell et al, 2016) and an analogous splicing variation (uniprot.org) between BMP2K and AAK1, some of the functions which we discovered for BMP2K variants could be possibly shared by AAK1 isoforms in other tissues.

**Figure 8.**
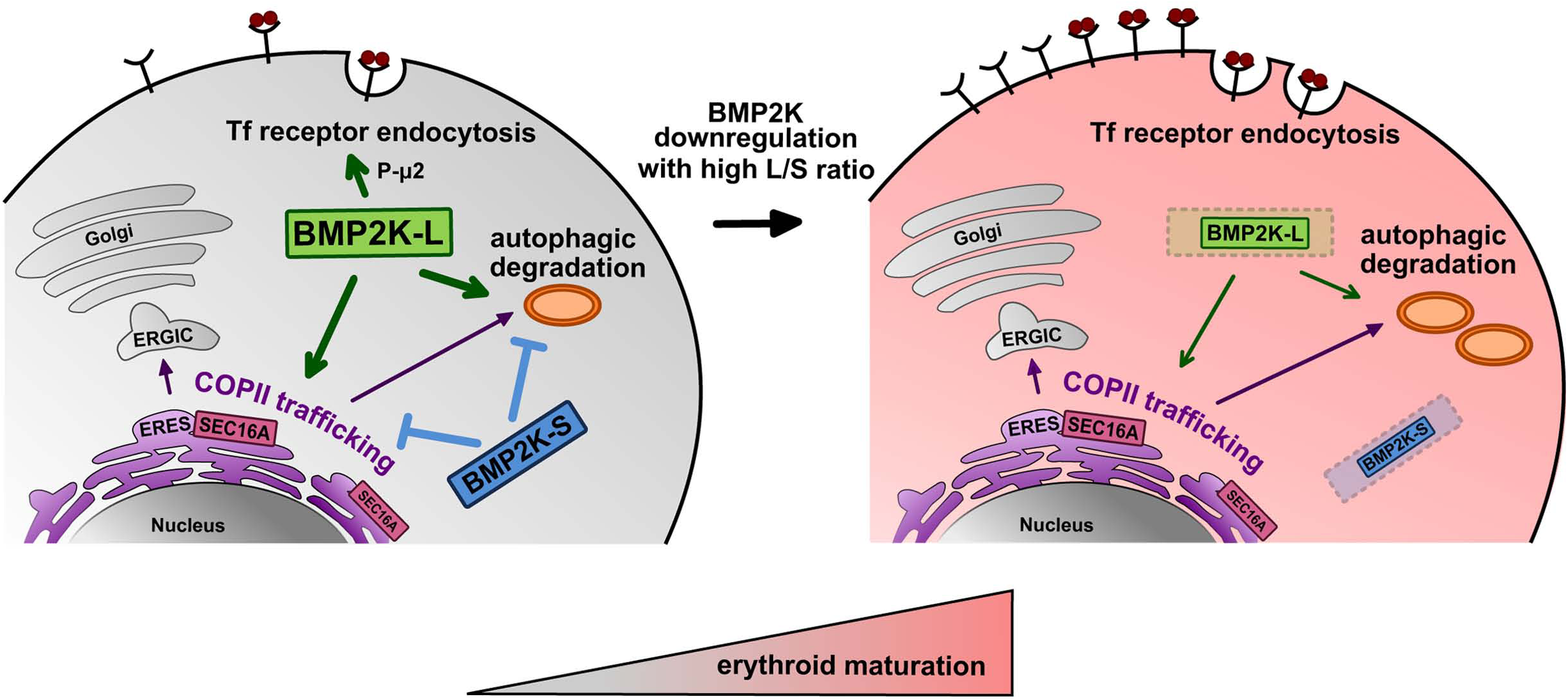
A model for the involvement of the BMP2K-L/S system in regulation of intracellular events important for erythroid maturation. BMP2K-L (the activator) and -S (the inhibitor) are both highly abundant in immature erythroblasts (left). Hence, in these cells the stimulating functions of BMP2K-L with respect to COPII trafficking and autophagy are inhibited by BMP2K-S. Upon downregulation of both BMP2K variants with preserved high L/S ratio, the inhibitory effects of BMP2K-S decline and the remaining BMP2K-L molecules stimulate COPII production at SEC16A-positive ERES and promote autophagy. These events facilitate erythroid maturation (right) associated with higher biosynthesis of transferrin (Tf) receptors. Higher abundance of Tf receptors at the PM fosters Tf uptake despite reduced Thr156 phosphorylation of μ2, and lower Tf CME efficiency due to BMP2K-L downregulation.

Enrichment of a protein during cell lineage differentiation may reflect its regulatory role, either positive or negative. BMP2K was proposed to negatively regulate osteoblast differentiation (Kearns et al, 2001) but to promote erythroid differentiation (Potts et al, 2013). We find that high levels of BMP2K variants restrict erythroid differentiation. However, erythroid maturation is associated not only with reduction of their levels but also with increased L/S ratio, i.e. the balance between variant levels shifted in favor of the longer (**Fig. 8**). We propose that BMP2K-L positively regulates cellular events required for erythroid maturation, while BMP2K-S inhibits these BMP2K-L-dependent functions. In such case, reducing the levels of BMP2K-S would favor pro-differentiation functions of BMP2K-L. However, as the two isoforms are splicing variants of one gene, they are transcriptionally co-regulated and therefore in cells neither of the variants should be present in very high excess. What mechanisms fine tune the L/S ratio during erythropoiesis remains to be studied.

Our data imply that the two BMP2K variants regulate intracellular events in an opposite manner. Based on proteomic reports (Borner et al, 2012; Brehme et al, 2009; Krieger et al, 2013) and high homology to AAK1, we initially assumed that BMP2K variants regulate endocytosis. Yeast homologues of BMP2K and AAK1 kinases, Akl1 and Prk1, suppress endocytosis (Bar-Yosef et al, 2018; Roelants et al, 2017; Takahashi et al, 2006; Zeng & Cai, 1999; Zeng et al, 2001). Consistently, we found that erythroid differentiation of K562 cells upon silencing of *BMP2K* expression correlated with elevated endocytic uptake. However, we did not find evidence for BMP2K variants acting as suppressors of CME. Conversely, we observed that BMP2K-L controls Thr156 phosphorylation of μ2 and promotes Tf endocytosis (**Fig. 8**). Moreover, we discovered that depletion of BMP2K isoforms with high L/S ratio must regulate cellular events other than Tf endocytosis to induce erythroid differentiation resulting in higher expression of genes encoding erythroid-enriched markers, such as Tf receptors. TFRC is a prototype receptor whose local clustering promotes CME (Liu et al, 2010). Thus, the elevated endocytic uptake upon *BMP2K* silencing could at least in part result from higher Tf receptor abundance, stimulating CME despite lower μ2 phosphorylation. However, we cannot exclude that either of the variants affect Tf uptake by direct or indirect regulation of bulk endocytosis, which may be induced upon CME inhibition (Kasprowicz et al, 2014).

We show that depletion of BMP2K variants strongly elevates hemoglobin production upon addition of hemin, a heme precursor which stimulates erythroid differentiation in a manner independent of Tf uptake (Fibach et al, 1987) but involving activation of autophagy (Grosso et al, 2019). Autophagic clearance of intracellular content is indispensable for erythroid differentiation (Cao et al, 2016; Grosso et al, 2017). We confirmed that, in addition to endocytosis, BMP2K isoforms regulate autophagy. Such role of BMP2K was proposed in a single study which did not analyze its underlying mechanisms or splicing variation of the kinase (Potts et al, 2013). In agreement with our proposal that the two BMP2K variants differentially regulate specific intracellular processes, we found BMP2K-L to promote autophagy while -S to inhibit autophagic flux (Fig. 8). Importantly, the induction of erythroid differentiation in K562 cells with reduced levels of both variants and high L/S ratio was associated with elevated autophagic flux. More efficient autophagic degradation in these cells may favor differentiation, as activated autophagy can by itself increase expression of genes encoding markers of erythroid maturation (Cao et al, 2016).

Although in the BioID interactomes we did not find any bona-fide autophagic regulators, we detected components of the ER-Golgi transport pathway that contributes to autophagosome formation (Ge et al, 2013; Shima et al, 2019; Wang et al, 2014). Among them was SEC16A, whose yeast homologue was shown as required for starvation-induced autophagy (Ishihara et al, 2001). Unexpectedly, we discovered that SEC16A in K562 cells is dispensable for basal autophagic degradation. Actually, in these cells the abundance of SEC16A protein inversely correlates with autophagic degradation: a) autophagic flux is increased in the absence of this protein, b) in cells lacking BMP2K-L, higher SEC16A protein levels are associated with autophagy inhibition, c) in cells lacking BMP2K-S, SEC16A level reduction coincides with elevated autophagic flux.

The involvement of SEC16A in autophagy is likely due to its role in COPII vesicle trafficking. We provide indications that the two possible functions described for yeast Sec16, i.e. organizing COPII assembly or inhibition of Sec31 recruitment (Glick, 2014; Sprangers & Rabouille, 2015) are performed by SEC16A in K562 cells, but in a manner dependent on its intracellular localization. We identified two distinct populations of COPII assembly sites. The juxtanuclear, where highly abundant SEC16A is required for efficient production of secretory COPII vesicles, and the dispersed, where less abundant SEC16A seems to restrict SEC31A recruitment, likely limiting the outer cage production. We propose that the SEC16A-dependent mechanisms at both sites are under control of the BMP2K-L/S system, in a manner coinciding with autophagy regulation. Specifically, higher recruitment of SEC31A to dispersed COPII vesicles was associated with elevated autophagic flux under two independent experimental conditions, i.e. upon SEC16A depletion or in cells lacking BMP2K-S. Similarly, TRAPPIII complex, one of COPII tethering factors that is known to promote autophagic flux (Behrends et al, 2010), was recently shown to positively modulate the recruitment of COPII outer cage (Zhao et al, 2017). However, whether there is a causal link between these two TRAPPIII functions remains unknown.

We also discover that the BMP2K-L/S-controlled COPII coating concerns vesicles positive for SEC24B (**Fig. 8**) shown to contribute to autophagy (Jeong et al, 2018). First, this reinforces the possible causality between BMP2K variant-mediated COPII trafficking and regulation of autophagy. Second, as in yeast the Sec16-dependent regulation of Sec31 recruitment is modulated by Sec24 (Kung et al, 2012) it is possible that either of the BMP2K variants cooperates with SEC24B in regulation of SEC16A function. Yeast Sec24 involvement in autophagy is modulated by multiple phosphorylation events (Davis et al, 2016). Hence, the BMP2K-L/S system may regulate COPII trafficking via phosphorylation of SEC16A or SEC24B or other COPII components. ERES-dependent trafficking is stimulated by phosphorylation of SEC16A by ULK1 (Joo et al, 2016) or ERK2 (Farhan et al, 2010). The phosphorylation by ULK1 occurs under basal conditions and does not affect autophagy, while that by ERK2 occurs upon growth factor signaling and its impact on autophagy has not been studied. Potentially, phosphorylation by BMP2K could be yet another modification to regulate SEC16A-dependent trafficking, possibly affecting autophagy. However, how such phosphorylation-dependent regulation by two counteracting splicing variants of a kinase would occur remains to be studied.

Our data suggest that, apart from autophagy, the BMP2K-L/S system regulates COPII-dependent secretion (**Fig. 8**). Modulation of SEC16A function in COPII coat recruitment affects the efficiency of secretion (Maeda et al, 2017; Wilhelmi et al, 2016). Higher SEC16A levels have been so far attributed to more efficient ERES-mediated trafficking (Tillmann et al, 2015). Conversely, we observed higher SEC16A abundance in cells with impaired COPII production due to absence of BMP2K-L. These cells also had reduced Tf receptor levels at the PM. It is thus possible that BMP2K-L positively regulates Tf receptor trafficking through ERES. On the other hand we show that BMP2K-S restricts recruitment of SEC31A to COPII vesicles. As studies in yeast suggest that Sec16-dependent inhibition of Sec31 recruitment is required for proper COPII turnover (Bharucha et al, 2013), strong SEC31A accumulation in BMP2K-S-depleted cells may reflect stalled COPII trafficking. Consistently, these cells show signs of strongly inhibited Tf receptor delivery to their PM. Nevertheless, reducing the levels of both variants but with preserved high L/S ratio seems to exert a balanced effect on COPII production and therefore efficient Tf receptor secretion, in response to its increased biosynthesis upon erythroid differentiation.

Endocytosis and autophagy (Fraser et al, 2017; Tooze et al, 2014) as well as secretory trafficking and autophagy (Davis et al, 2017; McCaughey & Stephens, 2018) are clearly interdependent. We provide evidence that an endocytic kinase regulates autophagy. This may occur at least in part via modulation of early secretory trafficking. Moreover, we find that BMP2K variants regulate endocytosis dependent on the AP-2 adaptor complex that has been shown to promote autophagosome formation and LC3B-mediated autophagic degradation (Popovic & Dikic, 2014; Tian et al, 2013). Hence, it is possible that the two BMP2K isoforms are cumulatively involved in the AP-2-dependent, SEC16A-dependent regulation of autophagy. Thus, through its alternative splicing variants, BMP2K kinase functions at the crossroad between endocytosis, secretion and autophagy. As tight coordination of these membrane trafficking pathways is particularly important for red blood cell maturation, therefore in the erythroid cell lineage *BMP2K* gene is highly expressed. Based on our discoveries, we propose that this gene encodes a regulatory system wherein the activator (BMP2K-L) promotes, while the inhibitor (BMP2K-S) restricts processes required for erythroid maturation. Hence, when the components of this system are highly expressed during erythropoiesis, reduced *BMP2K* gene expression promotes erythroid differentiation, provided that the amounts of the activator effectively surpass the levels of the inhibitor.

## MATERIALS AND METHODS

### Antibodies

The following antibodies were used: anti-BMP2K (sc-134284), anti-ARFGAP1 (sc-271303) and anti-GAPDH (sc-25778) from Santa Cruz; anti-phospho-μ2 (#7399), anti-REPS1 (#6404), anti-SEC24B (#12042) and anti-LC3B (#2775) from Cell Signaling Technologies; anti-EPS15R (ab76004) from Abcam; anti-TGN46 (T7576), anti-tubulin (T5168), anti-vinculin (V9131) and anti-beta-actin (A5441) from Sigma; anti-SQSTM1/p62 (610833), anti-μ2 (611350) and anti-SEC31A (612350) from BD Biosciences; anti-GFP (AF4240) from R&D Systems; anti-SEC16A (A300-648A) from Bethyl Laboratories Inc.; anti-SEC16A (HPA005684) from Atlas Antibodies; anti-TFRC (H68.4) from Thermo Fisher Scientific; secondary horseradish peroxidase (HRP)-conjugated goat anti-mouse and goat anti-rabbit antibodies from Jackson ImmunoResearch; secondary Alexa Fluor 488-conjugated anti-mouse, Alexa Fluor 555-conjugated anti-rabbit, and Alexa Fluor 647-conjugated anti-mouse antibodies from Thermo Fisher Scientific.

### Plasmids

To obtain pcDNA3.1-BMP2K-L construct, full-length human BMP2K-L was amplified from HEK293 cell cDNA by PCR using oligonucleotides 5’-ggggAAGCTTATGAAGAAGTTCTCTCGGATGCC-3’ (forward with HindIII restriction site) and 5’-ggggGGATCCCTACTGTTTAGAAGGAAATGGAGCAG-3’ (reverse with BamHI restriction site), subcloned into the pcDNA3.1 vector, and sequence-verified. To obtain pcDNA3.1-BMP2K-S construct, full-length human BMP2K-S was amplified from the clone IRAT32H09 (SourceBioScience) by PCR with the oligonucleotides 5’-ggggAAGCTTATGAAGAAGTTCTCTCGGATGCC-3’ (forward with HindIII restriction site) and 5’-AACAGCTATGACCATG-3’ (reverse M13 primer), subcloned into the pcDNA3.1 vector, and sequence-verified.

The pEGFP-BMP2K-L construct was obtained by restriction digestion of pcDNA3.1-BMP2K-L with HindIII and BamHI and subcloning the insert into pEGFP-C3 vector. The pEGFP-BMP2K-S construct was obtained by amplification of BMP2K-S from pcDNA3.1-BMP2K-S using oligonucleotides 5’-ggggAAGCTTATGAAGAAGTTCTCTCGGATGCC-3’ (forward with HindIII restriction site) and 5’-ggggGGATCCTTACTGTGAAGCAAAATAAGCCTTC-3’ (reverse with BamHI restriction site) and subcloning into pEGFP-C3 vector.

The pcDNA3.1-Myc-BirA*-BMP2K-L or pcDNA3.1-Myc-BirA*-BMP2K-S constructs were obtained by amplification of BMP2K-L or BMP2K-S from pcDNA3.1-BMP2K-L or pcDNA3.1-BMP2K-S using oligonucleotides 5’-gctaGGATCCTATGAAGAAGTTCTCTCGGAT-3’ (forward with BamHI restriction site) and 5’-gcgcGGTACCCTACTGTTTAGAAGGAAATG-3’ (reverse with KpnI restriction site for BMP2K-L) or 5’-gcgcGGTACCTTACTGTGAAGCAAAATAAG-3’ (reverse with KpnI restriction site for BMP2K-S) and subcloning into pcDNA3.1 mycBioID vector. The pcDNA3.1-BMP2K-L-BirA*-HA or pcDNA3.1-BMP2K-S-BirA*-HA constructs were obtained by amplification of BMP2K-L or BMP2K-S from pcDNA3.1-BMP2K-L or pcDNA3.1-BMP2K-S using oligonucleotides 5’-gcgcACCGGTATGAAGAAGTTCTCTCGGAT-3’ (forward with AgeI restriction site) and 5’-gcgcGGATCCCTGTTTAGAAGGAAATGGAG-3’ (reverse with BamHI restriction site for BMP2K-L) or 5’-gcgcGGATCCCTGTGAAGCAAAATAAGCCT-3’ (reverse with BamHI restriction site for BMP2K-S) and subcloning into pcDNA3.1 MCS-BirA(R118G)-HA vector. The pcDNA3.1 mycBioID and pcDNA3.1 MCS-BirA(R118G)-HA vectors (Addgene plasmids # 35700 and # 36047) were gifts from Kyle Roux (Roux et al, 2012).

BioGFP-μ2 construct, pUltra-Chili vector as well as psPAX2 and pMD2.G lentiviral packaging plasmids were a gift from J. Jaworski (International Institute of Molecular and Cell Biology in Warsaw, Poland). pEGFP-SEC16A construct (Addgene plasmid # 36155) was a gift from David Stephens (Watson et al, 2006). MISSION shRNA plasmids were obtained from Sigma-Aldrich. LentiCRISPRv2 vector was a gift from K. Mleczko-Sanecka (International Institute of Molecular and Cell Biology in Warsaw, Poland). gRNA sequences for CRISPR/Cas9 mediated gene inactivation were cloned into the LentiCRISPRv2 vector using a protocol described elsewhere (Sanjana et al, 2014). The pUltra-EGFP-BMP2K-L or pUltra-EGFP-BMP2K-S lentiviral constructs were obtained by digestion of pEGFP-BMP2K-L or pEGFP-BMP2K-S plasmids with AgeI and BamHI restriction enzymes and subcloning into pUltra-Chili vectors substituting their dTomato inserts with the EGFP-BMP2K coding sequences.

### Cell culture and treatment

HEK293 cells were maintained in Dulbecco’s modified Eagle’s medium (DMEM) and K562 cells in RPMI-1640 medium (Sigma-Aldrich). All media had high glucose concentration and were supplemented with 10% fetal bovine serum and 2 mM L-glutamine (Sigma-Aldrich). Hemin (H9039, Sigma-Aldrich) was used to stimulate erythroid differentiation of K562 cells at 20 μM concentration for 48 h or 72 h. Hemin 4 mM stocks were prepared according to the published protocol (Addya et al, 2004). Briefly, 13 mg of hemin was resuspended in 200 μl of 0.5 M sodium hydroxide, mixed with 250 μl of 1 M Tris (pH 7.8) and H2O was added to a final volume of 5 ml. K562 cells were also treated for 48 h with 75 nM Bafilomycin A1 (B1793, Sigma-Aldrich) to assess autophagic flux.

### Cell transfection and lentiviral transduction

For western blotting in HEK293 cells, 2.6*10^5^ cells were seeded per well in 12-well plates. For co-immunoprecipitation of proteins ectopically expressed in HEK293 cells, 6*10^5^ cells were seeded in 60 mm dishes. The cells were transfected after 24 h with plasmid DNA using Lipofectamine® 2000 Transfection Reagent (Thermo Fisher Scientific) according to the manufacturer’s protocol.

Lentiviral particles to transduce K562 cells were produced in HEK293T cells using psPAX2 and pMD2.G packaging plasmids as described elsewhere (Barde et al, 2010). For infection, 10*10^6^ K562 cells were grown in 10 ml of virus-containing RPMI-1460 medium for 24 h. To achieve shRNA-mediated depletion of BMP2K or SEC16A, MISSION shRNA plasmids were used. The empty pLKO.1 plasmid (SHC001), and the construct expressing non-targeting shRNA (SHC202) served as controls. After initial testing (not shown in the manuscript figures) of 5 different shRNA sequences (TRCN0000000914, TRCN0000000915, TRCN0000000916, TRCN0000000917, TRCN0000226438), only one (TRCN0000000915, shBMP2K) was found to efficiently downregulate BMP2K expression as assessed by western blotting. For SEC16A depletion, two shRNA sequences were used (TRCN0000246017, shSEC16A#1 and TRCN0000246019, shSEC16A#2). To achieve CRISPR/Cas9-mediated inactivation of genes of interest, for each gene, 4 different gRNA sequences (Doench et al, 2016) were cloned into the lentiCRISPRv2 or lentiCRISPRv2GFP vectors and were lentivirally introduced into K562 cells. Their efficiency of gene expression silencing was tested by western blotting and 2 sequences causing the strongest reduction of target protein levels were chosen for further experiments. gRNAs used in this study are listed in **Table S3**. For BMP2K depletion, cells were transduced with lentiviral particles containing one of two control (gCtr#1 and gCtr#2) or gene-targeting (gBMP2K#1, gBMP2K#2, gSEC16A#1 or gSEC16A#2) lentiCRISPRv2 vectors for 24 h and selected with 2 µg/ml puromycin for 72 h.

For overexpression of EGFP-tagged BMP2K variants in K562 cells, cells were transduced for 24 h with lentiviral particles containing pUltra-EGFP-BMP2K-L or -S plasmids and analyzed after 3 subsequent days of culture.

### Mouse fetal liver erythroblast isolation, *ex vivo* differentiation and FACS

Mouse fetal liver erythroid progenitors were isolated and stimulated for differentiation as described (Zhao et al, 2014). Briefly 8-10 livers were extracted from embryonic day 13.5 C56/BL6 mouse embryos and manually disrupted. Ter119-positive differentiated cells were recognized using specific biotin-conjugated antibodies (13-5921-81) and removed from the cell suspension using Streptavidin-coupled magnetic beads (65001; both reagents from Thermo Fisher Scientific). To induce erythroid differentiation, 4*10^5^ purified and washed cells were cultured on fibronectin-coated 12-well plates in the presence of erythropoietin (EPO)-containing Iscove’s Modified Dulbecco’s Medium: 2 U/ml EPO (11120166001), 10 μg/ml insulin (I3536), 200 μg/ml holo-transferrin (T0665), 0.1 mM β-mercaptoethanol (M3148), 1% BSA, 15% FBS and 2 mM L-glutamine (all reagents from Sigma-Aldrich). After 24 h, 48 h or 72 h of differentiation the cells were harvested for western blotting or qRT-PCR analyses.

To analyze protein levels in mouse erythroblasts at various stages of differentiation, the isolated mouse fetal liver progenitors were differentiated for 96 h and labeled with APC-conjugated anti-Tfrc and PE-conjugated anti-Ter-119 antibodies as described (Zhao et al, 2014). The labeled cells were FACS-separated using BD FACSAria II cell sorter and harvested for western blotting analysis.

### Immunofluorescence staining and microscopy

K562 cells (non-treated or transduced with lentiviral plasmids) were transferred to ice, fixed with 3% paraformaldehyde for 15 min on ice followed by 15 min at room temperature. After three washes with PBS, cells were immunostained in suspension using an adapted version of protocol described for adherent cells (Maminska et al, 2016). In most cases, stained cells were resuspended in 0.5% low-melting agarose (Sigma-Aldrich) and transferred to microscopy 96-well plates (Greiner Bio-One). The plates were scanned using Opera Phenix high content screening microscope (PerkinElmer) with 40×1.1 NA water immersion objective. Harmony 4.9 software (PerkinElmer) was used for image acquisition and their quantitative analysis. To quantify chosen parameters in the obtained images (mean pixel intensity, number of vesicles, mean fluorescence intensity per vesicle as well as Pearson’s coefficient), more than 40 microscopic fields were analyzed per each experimental condition. Non-treated K562 cells stained for ARFGAP1 were mounted using ProLong® Gold Antifade Reagent (Thermo Fisher Scientific) on glass slides. The slides were scanned using a ZEISS LSM 800 confocal microscope with a 40×1.3 NA oil immersion objective. ZEN Blue 2.3 (Version 2.3.69.1003) (Zeiss) was used for image acquisition. Pictures were assembled in Photoshop (Adobe) with only linear adjustments of contrast and brightness.

### Western blotting

Cells were lysed in RIPA buffer (1% IGEPAL® CA-630, 0.5% sodium deoxycholate, 0.1% SDS, 50 mM Tris (pH 7.4), 150 mM NaCl, 0.5 mM EDTA) supplemented with protease inhibitor cocktail (6 μg/ml chymostatin, 0.5 μg/ml leupeptin, 10 μg/ml antipain, 2 μg/ml aprotinin, 0.7 μg/ml pepstatin A and 10 μg/ml 4-amidinophenylmethanesulfonyl fluoride hydrochloride; Sigma-Aldrich) and phosphatase inhibitor cocktails (P0044 and P5726, Sigma-Aldrich). For HEK293 or K562 cells, protein concentration was measured with BCA Protein Assay Kit (Thermo Scientific) and 10-50 µg of total protein per sample were resolved on SDS-PAGE. For mouse fetal liver erythroblasts, RIPA lysate equivalents of 3*10^5^ cells were resolved on SDS-PAGE. Resolved proteins were transferred to nitrocellulose membrane (Whatman), probed with specific primary and secondary antibodies, and detected using ChemiDoc imaging system (Bio-Rad) or Odyssey infrared imaging system (LI-COR Biosciences).

Densitometric analysis of western blotting bands was performed using Image Lab 5.2.1 software (Bio-Rad). The raw data were normalized to tubulin band intensities (for K562 cells) or were not normalized (for mouse fetal liver erythroblasts) and presented as fold levels over respective controls.

### Co-immunoprecipitation (Co-IP)

For co-immunoprecipitation of proteins ectopically expressed in HEK293 cells or endogenous proteins in K562 cells, an equivalent of 8*10^5^ cells in 300 μl of RIPA buffer was used per reaction. The lysates were incubated for 1.5 h at 4°C with 1 µg of specific antibodies (goat anti-GFP, rabbit anti-REPS1, rabbit anti-phospho-μ2, rabbit anti-SEC24B or unspecific rabbit IgG) with rotation. The immune complexes were captured by incubation with Protein G-agarose for 2 h at 4°C with rotation. The agarose beads-bound protein complexes were spun down and washed three times using IP wash buffer (50 mM Hepes, pH 7.5, 300 mM NaCl, 1 mM EGTA, 1 mM EDTA, 1% Triton X-100, 10% glycerol, 5 μg/ml DNase and protease inhibitor cocktail) and one time using 50 mM Hepes, pH 7.5. The proteins were eluted from the beads by incubation at 95°C for 10 min with Laemmli buffer and analyzed by western blotting. Protein G-agarose beads were blocked in PBS with 5% BSA for 2 h at 4°C with rotation before they were added to capture the immune complexes.

### Quantitative real-time PCR (qRT-PCR)

Total RNA was isolated from cells with High Pure Isolation Kit (Roche). For cDNA synthesis random nonamers, oligo(dT)23 and M-MLV reverse transcriptase (Sigma-Aldrich) were used according to the manufacturer’s instructions. To estimate the expression of genes of interest we used primers designed with NCBI tool (and custom-synthesized by Sigma-Aldrich) listed in **Table S4**. The qRT-PCR reaction was performed with the Kapa Sybr Fast ABI Prism qPCR Kit (KapaBiosystems) using a 7900HT Fast Real-Time PCR thermocycler (Applied Biosystems) with at least three technical repeats per experimental condition. The data were normalized according to the expression level of housekeeping gene, *GAPDH* (in K562 cells) or *Rpl19* (in mouse fetal liver erythroblasts) and presented as fold changes.

### Transferrin and dextran uptake assay

For continuous transferrin (Tf) uptake experiments, 1*10^6^ K562 cells were pre-incubated for 30 min on ice with 25 μg/ml Alexa Fluor 647-conjugated Tf (Thermo Fisher Scientific) and incubated at 37°C for 5 or 40 min (early uptake or steady-state loading, respectively) followed by washing and fixation with 3% PFA for 5 min. For dextran uptake measurement, cells were incubated with 125 μg/ml Alexa Fluor 488-conjugated 10 kDa dextran (Thermo Fisher Scientific) at 37°C for 40 min followed by washing and fixation.

The pulse-chase Tf uptake was performed according to a described protocol (Dannhauser et al, 2017) but adapted for K562 cells cultured in suspension. Briefly, non-starved cells were incubated for 15 min on ice with 5 μg/ml Alexa Fluor 647-conjugated Tf (Thermo Fisher Scientific) in RPMI medium, followed by washing unbound Tf with ice-cold PBS. Then cells were either analyzed immediately by flow cytometry, to assess the cell surface Tf binding, or were resuspended in pre-warmed RPMI medium and incubated at 37°C for 5 min to initiate Tf uptake. The uptake was stopped by washing in ice-cold PBS and non-internalized Tf was stripped by two acid washes (0.15 m glycine buffer, pH 3). Upon final wash with ice-cold PBS, cells were analyzed by flow cytometry. For each experimental conditions we applied control conditions, including non-stained cells (without Alexa Fluor 647-conjugated Tf incubation to estimate background fluorescence), and cells stripped immediately upon incubation with Tf (to determine the efficiency of surface-bound Tf removal).

Fixed (continuous uptake) or non-fixed (pulse-chase uptake) cells were resuspended in PBS and the fluorescence of incorporated Tf or dextran was recorded from 50 000 cells with a BD LSR Fortessa flow cytometer. The data collection and calculation of mean fluorescence intensities was performed using BD FACSDiva 6.2 software. Flow cytometry data were analyzed and visualized by FlowJo software (BD Biosciences).

### Hemoglobin content analysis

Hemoglobin content in K562 cells was visualized by staining with benzidine dihydrochloride (B3383, Sigma-Aldrich). Briefly, 300 µl of 2 mg/ml solution in 3% acetic acid was added to an equal volume of fresh RPMI-1640 medium with 2*10^5^ cells, followed immediately by addition of 12 µl of 30% hydrogen peroxide solution. The stained cells were imaged in bright-field using the IX 7C Olympus microscope and the percentage of hemoglobin-positive cells was counted using the ImageJ software. At least 1000 cells were counted per sample.

### Proximity biotinylation (BioID)

The expression of Myc-BirA*-BMP2K-L, Myc-BirA*-BMP2K-S, BMP2K-L-BirA*-HA or BMP2K-S-BirA*-HA fusion proteins was assessed after transient transfection of HEK293 cells. The obtained constructs as well as empty vectors were linearized using PvuI restriction enzyme and introduced into HEK293 cells using Lipofectamine® 2000 Transfection Reagent. The cells underwent antibiotic selection with 500 μg/ml G418 (Gibco) and clones stably expressing each of the fusion proteins were generated. The expression efficiencies among the obtained clones were assessed by western blotting (not shown in the manuscript figures) and two clones expressing the highest levels of each fusion protein were selected for biotin labelling. Biotin labelling and subsequent pull down of biotinylated proteins using NanoLink^(tm)^ Streptavidin Magnetic Beads, 0.8 µm (VWR) was performed in two biological repeats for each of the two selected clones, following the originally described protocol (Roux et al, 2013). The qualitative analysis of biotinylated proteins from each sample was performed by mass spectrometry and analyzed using Mascot software. The Mascot scores obtained for each of the two clones per condition (Myc-BirA*-BMP2K-L, Myc-BirA*-BMP2K-S, BMP2K-L-BirA*-HA or BMP2K-S-BirA*-HA) were averaged and the scores obtained for proteins identified in the control samples (false-positive background of Myc-BirA* or BirA*-HA) were subtracted. The final subtracted C-tag or N-tag scores were obtained from averaging the subtracted scores from two biological repeats.

### Mass spectrometry of biotinylated proteins

Proteins biotinylated in BioID were reduced with 5 mM TCEP (for 60 minutes at 60°C). To block reduced cysteines, 200 mM MMTS at a final concentration of 10 mM was added and the sample was incubated at room temperature for 10 minutes. Trypsin (Promega) was added at a 1:20 vol./vol. ratio and incubated at 37°C overnight. Finally, trifluoroacetic acid was used to inactivate trypsin. Peptide mixtures were analyzed by liquid chromatography coupled to tandem mass spectrometry (LC-MS/MS) using Nano-Acquity (Waters Corporation) UPLC system and LTQ-FT-Orbitrap (Thermo Scientific) mass spectrometer. Measurements were carried out in the positive polarity mode, with capillary voltage set to 2.5 kV. A sample was first applied to the Nano-ACQUITY UPLC Trapping Column using water containing 0.1% formic acid as a mobile phase. Next, the peptide mixture was transferred to Nano-ACQUITY UPLC BEH C18 Column using an acetonitrile gradient (5–35% acetonitrile over 160 minutes) in the presence of 0.1% formic acid with a flow rate of 250 nl/min. Peptides were eluted directly to the ion source of the mass spectrometer. Each LC run was preceded by a blank run to ensure that there was no carry-over of material from previous analysis. HCD fragmentation was used. Up to 10 MS/MS events were allowed per each MS scan.

Acquired raw data were processed by Mascot Distiller followed by Mascot Search (Matrix Science, London, UK, on-site license) against SwissProt database restricted to human sequences. Search parameters for precursor and product ions mass tolerance were 30 ppm and 0.1 Da, respectively, enzyme specificity: trypsin, missed cleavage sites allowed: 1, fixed modification of cysteine by methylthio and variable modification of methionine oxidation. Peptides with Mascot score exceeding the threshold value corresponding to <5% expectation value, calculated by Mascot procedure, were considered to be positively identified. BioID-MS detection scores were visualized using R package ggplot2 and scales (R version 3.4.4).

The mass spectrometry proteomics data have been deposited to the ProteomeXchange Consortium via the PRIDE (Perez-Riverol et al, 2019) partner repository with the dataset identifier PXD013542.

### *In silico* protein amino acid sequence analysis

To identify amino acid motifs in the sequences of BMP2K splicing variants the eukaryotic linear motif (ELM) resource - 2018 update was used (Gouw et al, 2018).

### Statistical analysis

Data are provided as mean ± SEM. Statistical analysis was performed with Prism 6 (GraphPad Software) using unpaired two-tailed Student t test (for qRT-PCR analysis and western blotting densitometry in K562 cells) or paired two-tailed Student t test (for uptake experiments, hemoglobin content analysis, western blotting densitometry in mouse fetal erythroblasts and quantified parameters from confocal microcopy analysis). Data points were marked according to the p value, where p>0.05 is left unmarked or indicated with p=value, *p<0.05, **p<0.01, ***p<0.001, ****<0.0001.

## ACKNOWLEDGEMENTS

We are grateful to J. Jaworski and K. Mleczko-Sanecka for providing plasmids. We also thank M. Banach-Orłowska, M. Maksymowicz, A. Poświata, L. Wolińska-Nizioł and D. Zdżalik-Bielecka for critical reading of the manuscript. This work was funded by the MAESTRO grant (UMO-2011/02/A/NZ3/00149) from National Science Center to M. Miaczynska. M. Miaczynska, M. Kaczmarek and K. Jastrzębski were supported by TEAM grant (POIR.04.04.00-00-20CE/16-00), J. Cendrowski and M. Mazur supported by HOMING grant (POIR.04.04.00-00-1C54/16-00), K. Piwocka was supported by TEAM-TECH Core Facility Plus grant (POIR.04.04.00-00-23C2/17-00) – the three grants from the Foundation for Polish Science co-financed by the European Union under the European Regional Development Fund. Mass spectrometric analysis was performed in the Mass Spectrometry Laboratory, IBB PAS, Warsaw. The MS equipment was sponsored in part by the Centre for Preclinical Research and Technology (CePT), a project co-sponsored by European Regional Development Fund and Innovative Economy, The National Cohesion Strategy of Poland.

## AUTHOR CONTRIBUTIONS

The research was conceived by J.C. and M. Miaczynska. Funding was acquired by M. Miaczynska. Experiments were designed and performed mostly by J.C. with support from M.K., K.K-K., M. Mazur and K.J. and crucial help from K.P., M.B.O. and A.K. (flow cytometry). The manuscript was written by J.C. and M. Miaczynska. M. Miaczynska supervised the work. All authors approved the manuscript.

## SUPPLEMENTARY FIGURE LEGENDS

**Cendrowski et al. supplementary figures and tables**

**Figure S1.**
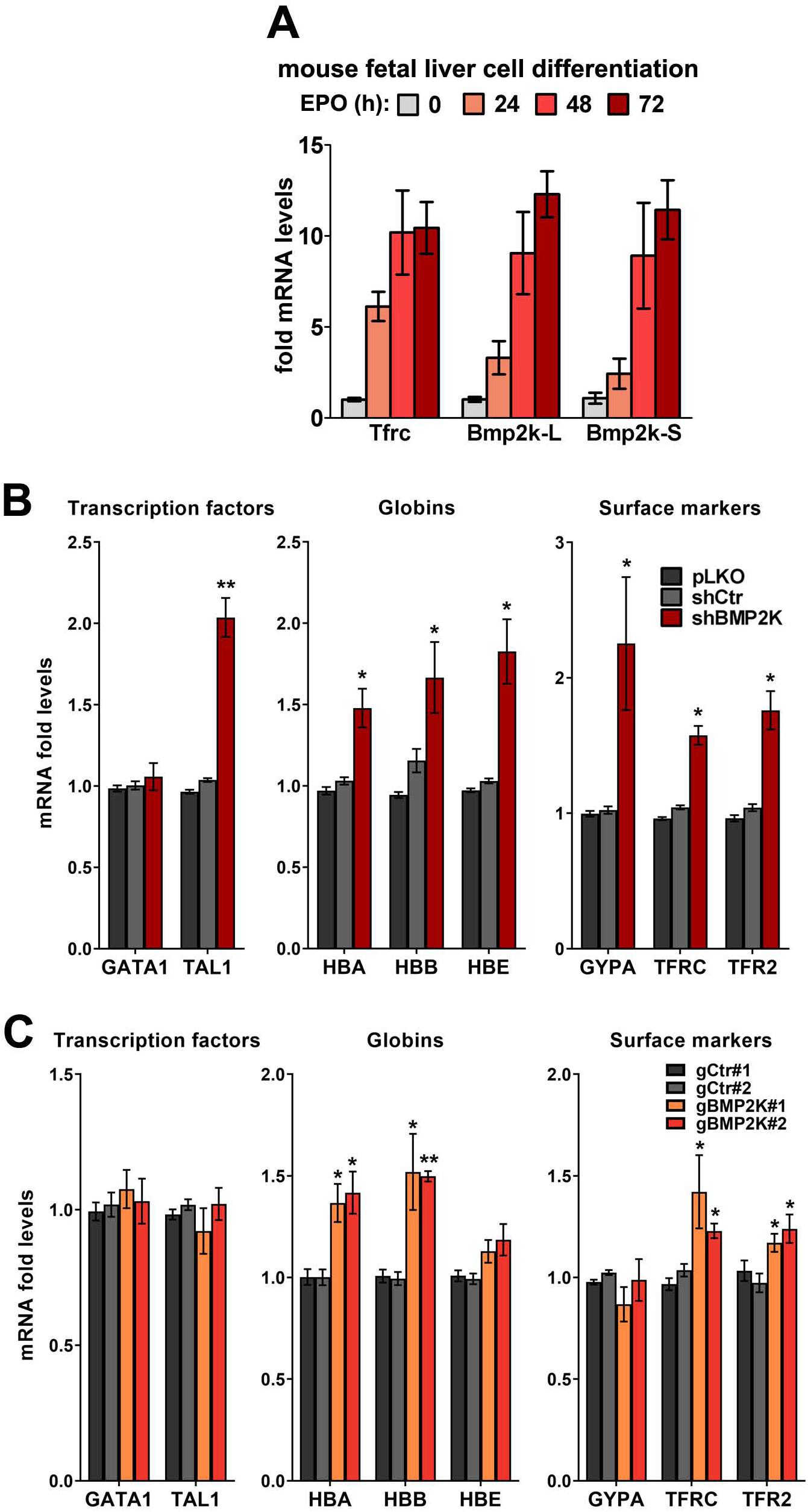
**A.** Fold changes in mRNA levels of transferrin receptor 1 (Tfrc) as well as of the longer or shorter Bmp2k splicing variants in erythroblasts isolated from mouse fetal liver and differentiated for the indicated time periods using erythropoietin (EPO)-containing medium (n=4 +/- SEM). **B and C.** Fold changes in mRNA levels of the indicated erythroid markers in control cells or in cells depleted of all BMP2K splicing variants using shRNA (B) or CRISPR/Cas9 (C) approach (n=3 or 4 in B and 3 in C +/- SEM). Control constructs used were: empty pLKO vector and non-targeting shRNA, shCtr (B) or non-targeting gRNAs, gCtr#1 and #2 (C). Fold changes in BMP2K-depleted cells were compared statistically to those in shCtr-treated cells (B) or to the average of values measured for gCtr#1 and gCtr#2-transduced cells (C). *p<0.05, **p<0.01.

**Figure S2.**
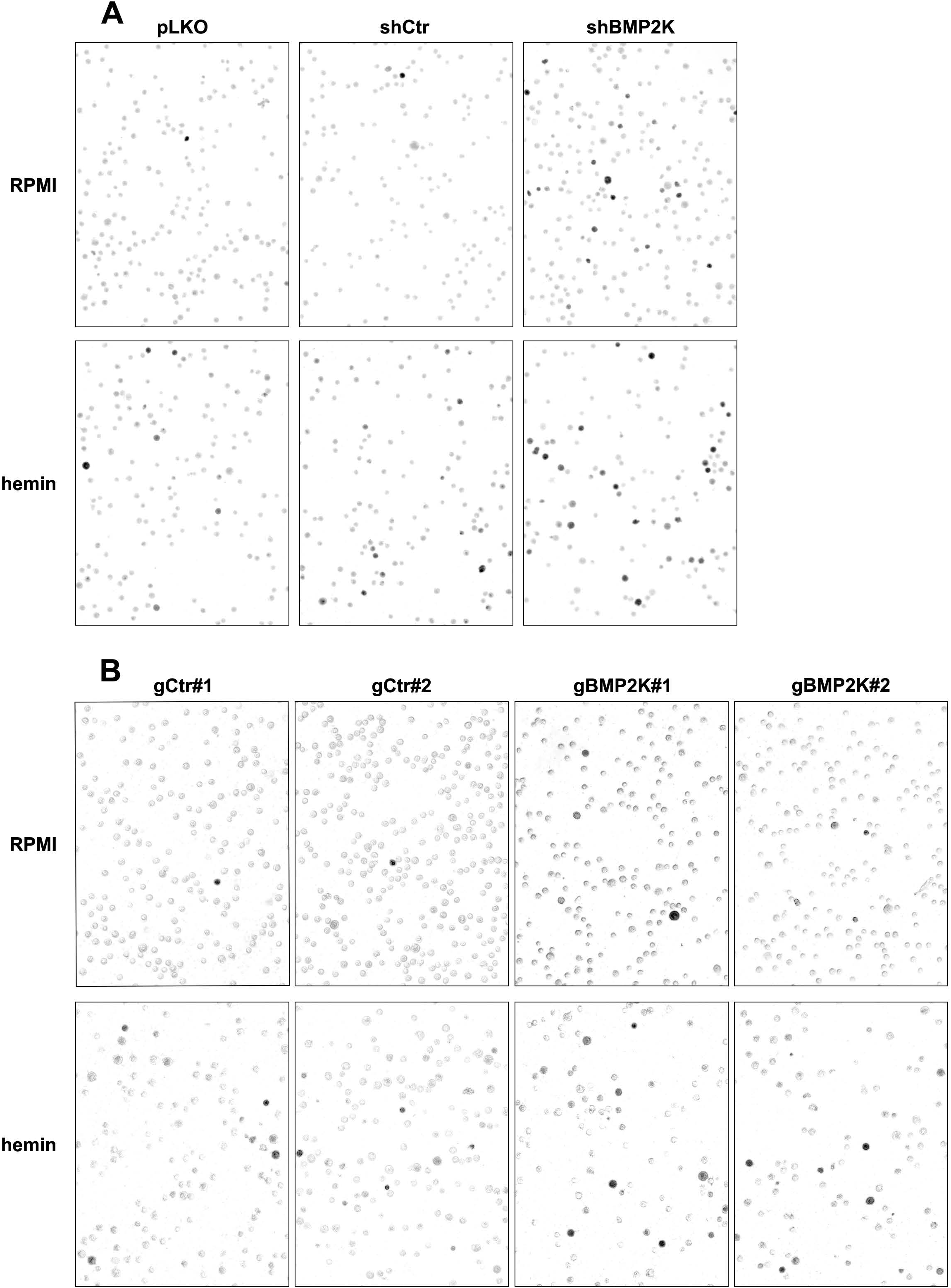
**A and B.** Representative images showing benzidine staining of non-treated (RPMI medium) or 20 µM hemin-treated (48 h) K562 cells, control or depleted of all BMP2K splicing variants using shRNA (A) or CRISPR/Cas9 (B) approach. Dark grey/black cells contain hemoglobin.

**Figure S3.**
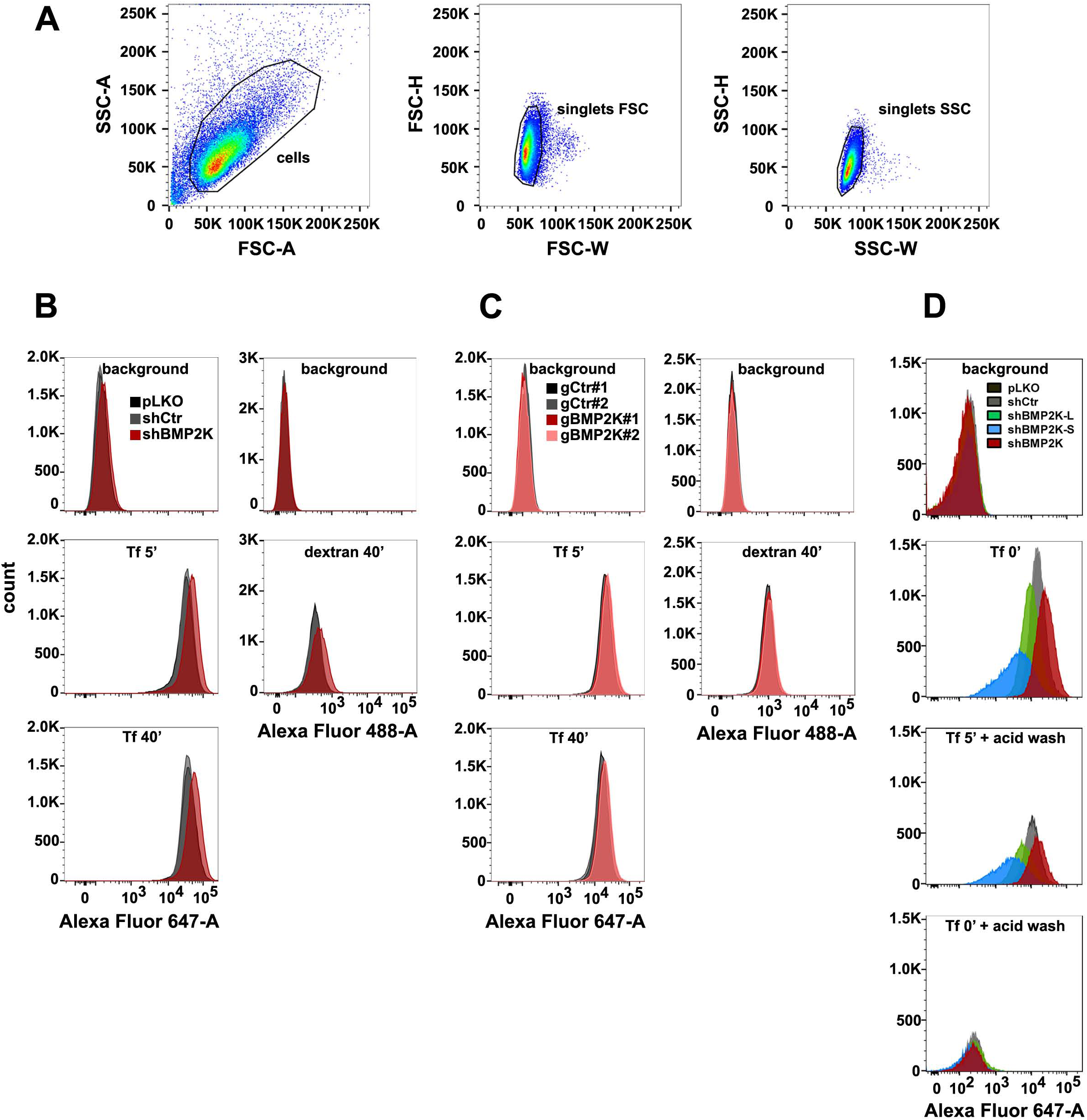
**A.** An example of gating strategy for flow cytometric analyses shown in Fig. 2 and 3 applied for K562 cells transduced with empty pLKO vector, including cell gate (FSC vs SSC) and singlet discrimination gates (FSC-H vs FSC-W, SSC-H vs SSC-W). This strategy was applied for all flow cytometry experiments. **B and C.** Overlay histograms showing fluorescence signal of AlexaFluor-647 or AlexaFluor-488, indicating Tf or dextran abundance respectively, in control cells or cells depleted of *BMP2K* gene products using shRNA (B) or CRISPR/Cas9 (C) approach. Control constructs used were: empty pLKO vector and non-targeting shRNA, shCtr (B) or non-targeting gRNAs, gCtr#1 and #2 (C). **D.** Overlay histograms showing fluorescence signal of AlexaFluor-647, indicating Tf abundance during pulse-chase endocytosis assay in control cells or cells depleted of BMP2K splicing variants. The histograms (B, C and D) illustrate representative experiments included in the analyses shown in Fig. 2 and 3.

**Figure S4.**
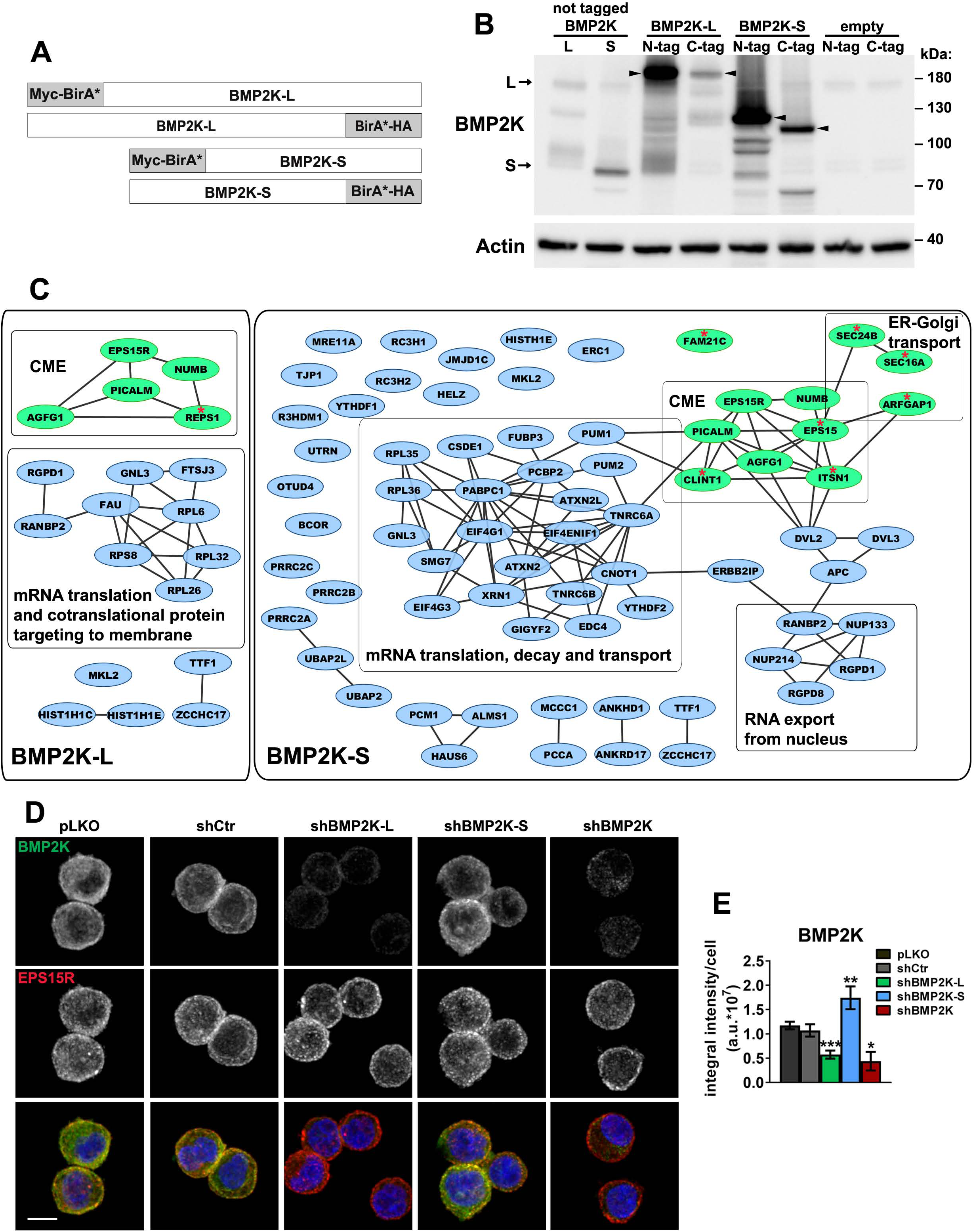
**A.** Schematic representation of fusion proteins that were ectopically expressed in HEK293 cells for the purpose of the proximity biotinylation (BioID) analysis. **B.** Western blots showing transient expression in HEK293 cells of BMP2K-L or BMP2K-S, N- or C-terminally tagged with BirA* biotin ligase as well as non-tagged L or S isoforms. Bands representing the full-length proteins are indicated by arrows (for non-tagged L and S isoforms) or arrowheads (for BirA*-tagged proteins). Actin was used as a gel loading control. **C.** Cytoscape diagrams (www.cytoscape.org) showing interactions between proteins detected in BioID as proximal to BMP2K-L or -S isoforms with indications of functionally related protein groups (CME – clathrin-mediated endocytosis). Green color indicates known regulators of vesicular trafficking. Red asterisks mark vesicular trafficking-related proteins detected as proximal to only one of the isoforms. **D.** Representative maximum intensity projection images from confocal microscope showing intracellular localization of endogenous BMP2K with respect to endogenous EPS15R in K562 cells depleted of single (shBMP2K-L or shBMP2K-S) or all BMP2K splicing variants (shBMP2K). Cell nuclei marked with DAPI stain (blue). Scale bar, 10 µm. **E.** Integral fluorescence intensity of BMP2K staining, shown as arbitrary units (a.u.), quantified from microscopic images of cells depleted of BMP2K splicing variants as represented by those in D (n=4 +/- SEM). Values of BMP2K integral intensity in cells depleted of BMP2K variants were compared statistically to the levels in cells treated with shCtr. *p<0.05, **p<0.01, ***p<0.01.

**Figure S5.**
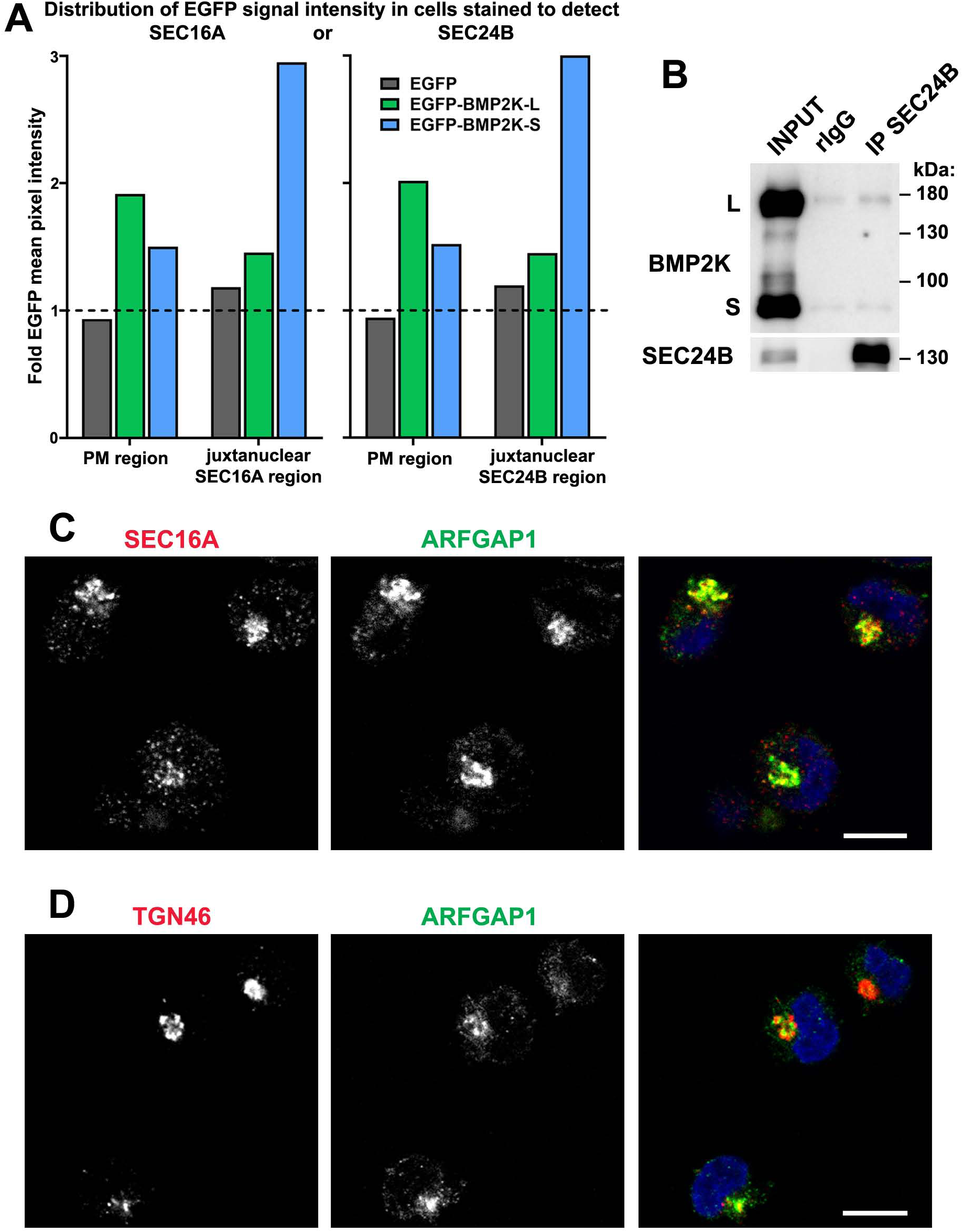
**A.** Mean pixel intensity of EGFP fluorescence signal in the indicated cell regions shown as fold change over mean intensity in the remaining cell area (represented by dashed horizontal line). Quantification performed based on images of cells represented in Fig. 4B: transduced with EGFP only, or EGFP-tagged BMP2K variants and stained using antibodies detecting SEC16A or SEC24B proteins (n=1, >4000 cells/condition). **B**. Western blots showing the levels of endogenous BMP2K and SEC24B in K562 cell lysates (INPUT) or samples after immunoprecipitation (IP) using either control immunoglobulins (rIgG) or antibodies recognizing SEC24B protein. **C and D.** Single plane confocal microscopy images showing localization of SEC16A (C) or TGN46 (D) with respect to ARFGAP1 in K562 cells. Scale bars, 10 µm.

**Figure S6.**
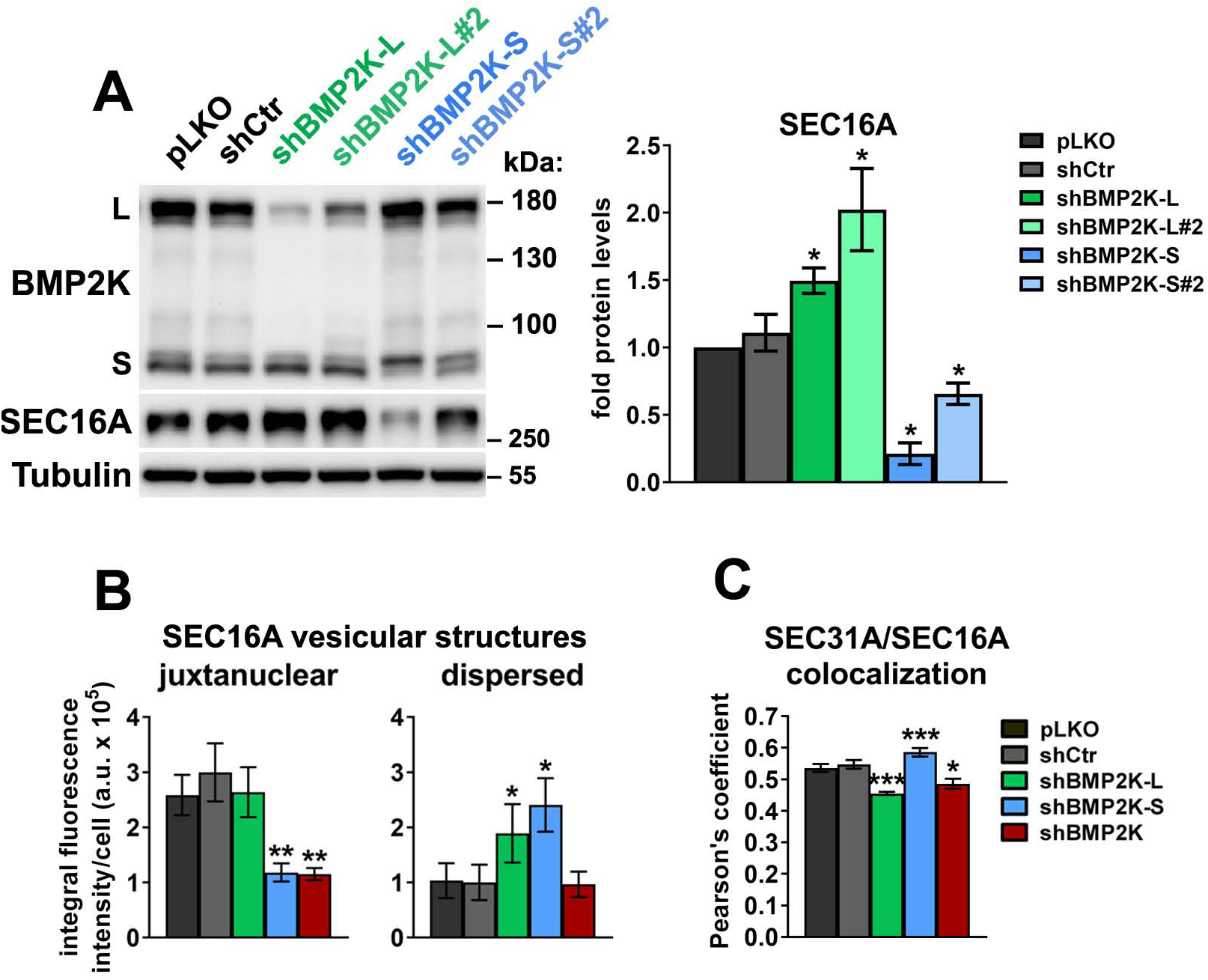
**A.** Western blots showing the efficiencies of BMP2K-L or -S depletion by additional shRNAs (shBMP2K-L#2 and shBMP2K-S#2) and their effects on SEC16A protein levels as compared to control vectors (pLKO, shCtr) and shRNAs shown in Fig. 6 (shBMP2K-L and shBMP2K-S). Graph shows densitometric analysis of western blotting bands for the indicated proteins using tubulin abundance for normalization (n=3 +/- SEM). **B and C.** Integral fluorescence intensity of juxtanuclear or dispersed SEC16A vesicles presented in arbitrary units (a.u.) per cell (B) and total colocalization between SEC31A and SEC24B structures quantified using the Pearson’s correlation coefficient (C) in control cells or in cells depleted of single (shBMP2K-L or shBMP2K-S) or all BMP2K splicing variants (shBMP2K). Quantifications performed based on images represented by those shown in Fig. 6C (n=5 +/- SEM). Values in cells depleted of BMP2K variants (A-C) were compared statistically to the levels in cells treated with shCtr. *p<0.05, **p<0.01, ***p<0.01.

**Figure S7.**
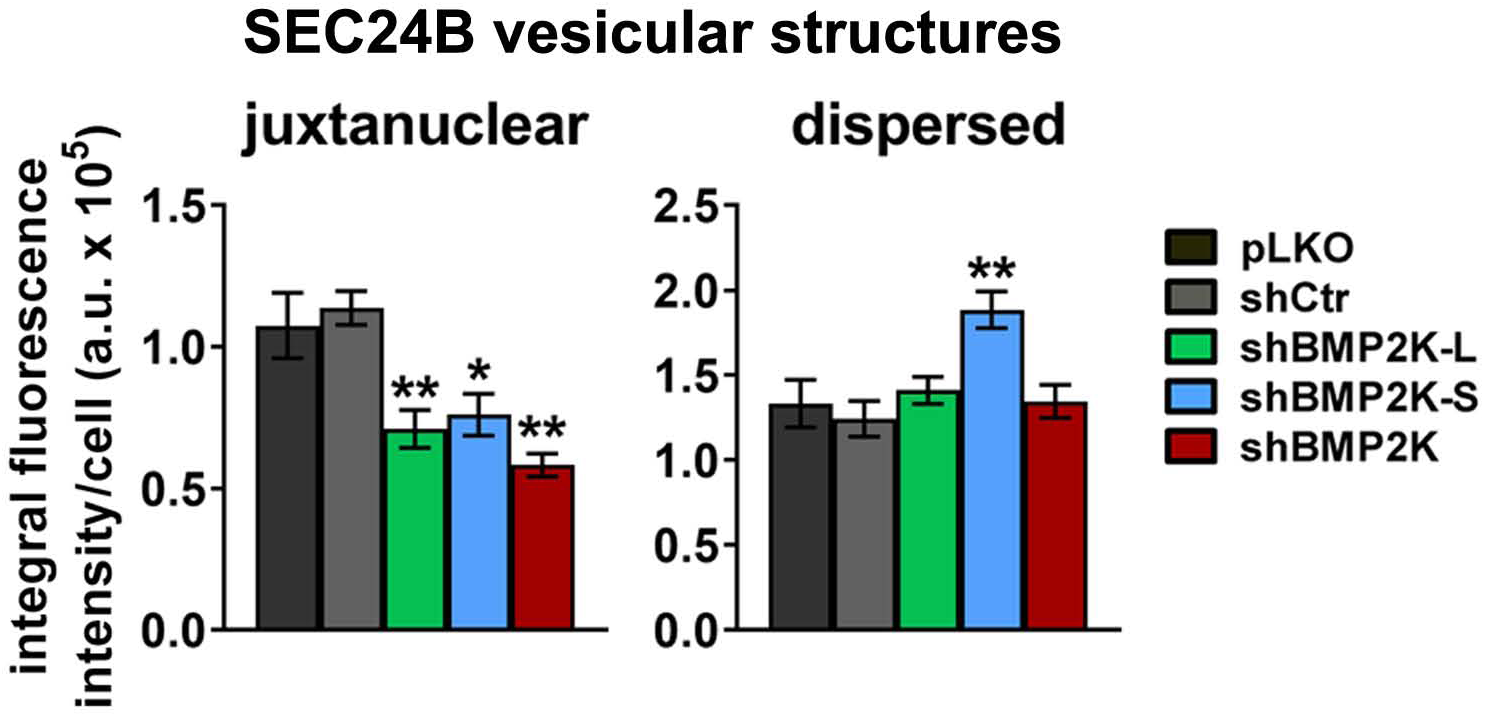
Integral fluorescence intensity of juxtanuclear or dispersed SEC24B vesicles presented in arbitrary units (a.u.) per cell, in cells depleted of single (shBMP2K-L or shBMP2K-S) or all BMP2K splicing variants (shBMP2K) as compared to control cells (pLKO, shCtr). Quantifications performed based on images represented by those shown in Fig. 7A (n=4 +/- SEM). Values in cells depleted of BMP2K variants were compared statistically to the levels in cells treated with shCtr. *p<0.05, **p<0.01.

## SUPPLEMENTARY TABLES

**Table S1.**
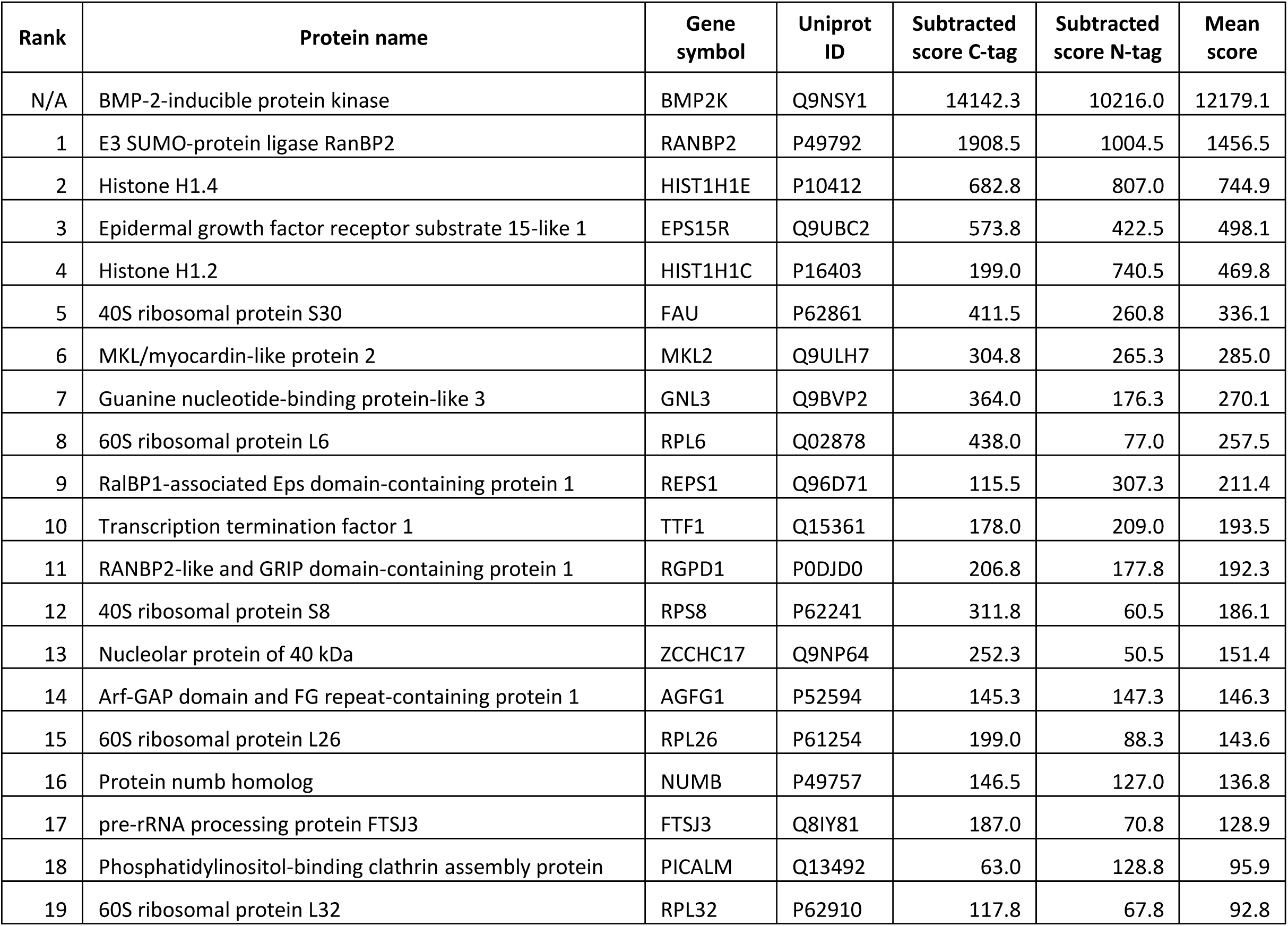
The ranked list of proteins detected in BioID using BMP2K-L as bait.

**Table S2.**
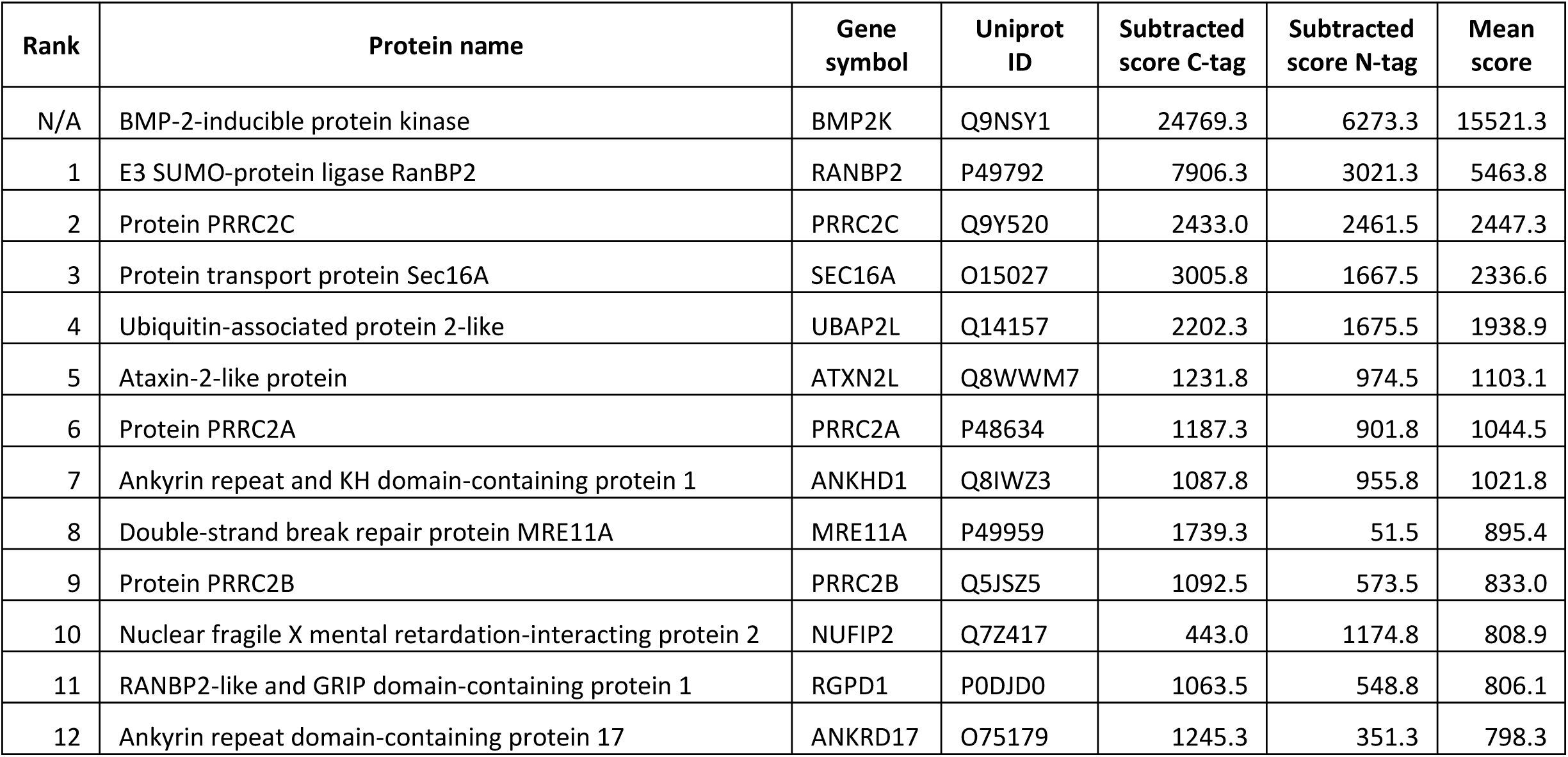
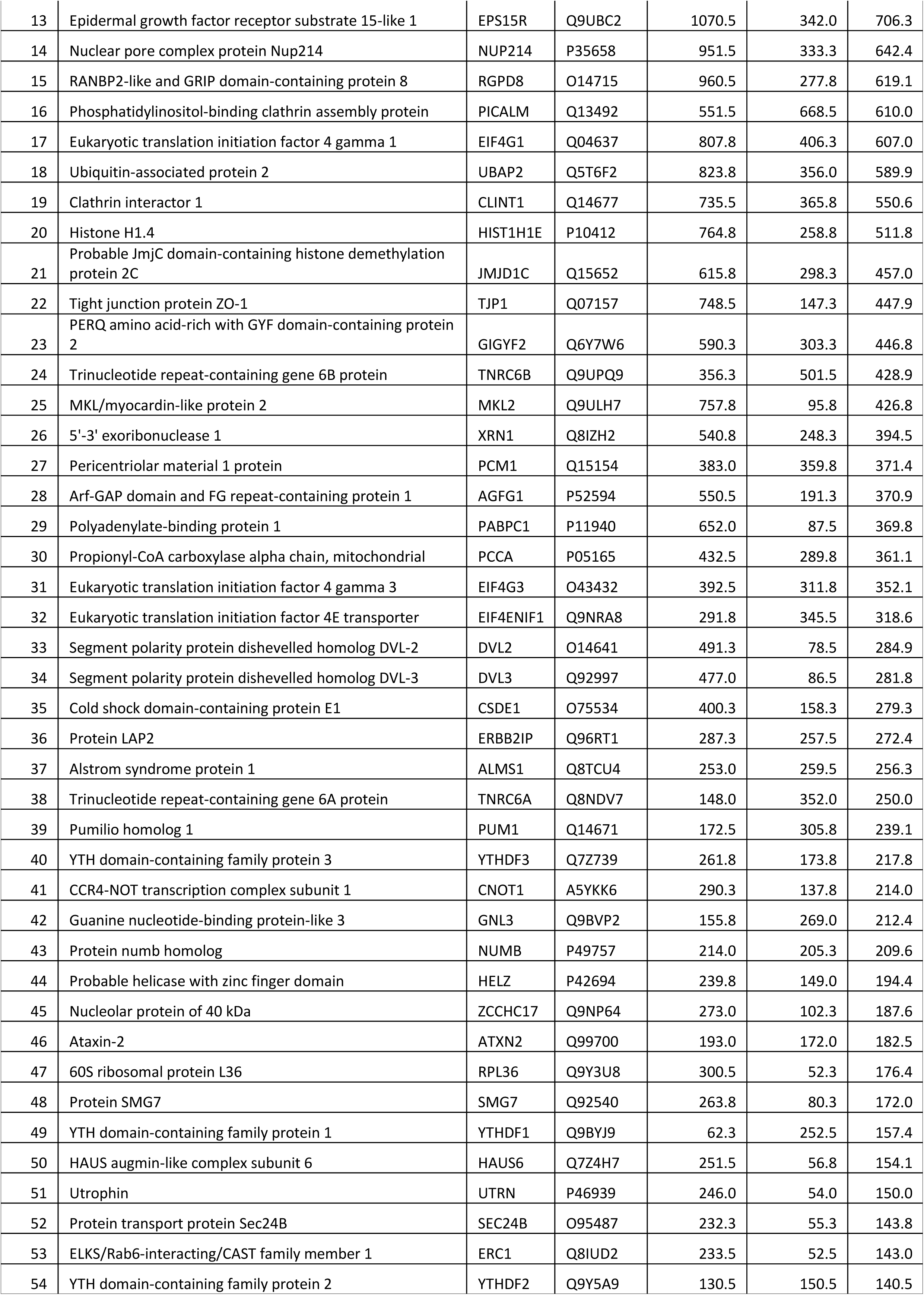
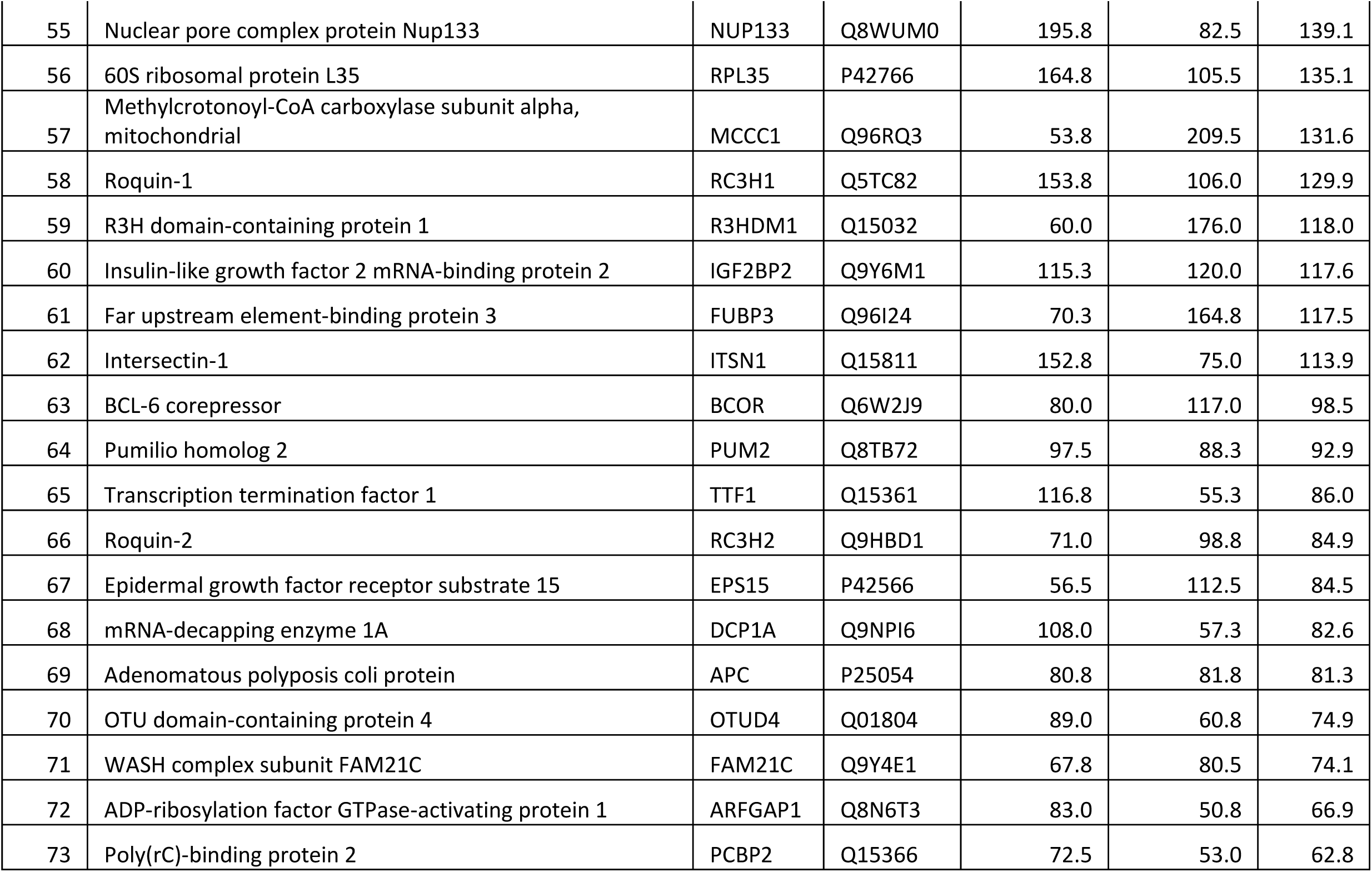
The ranked list of proteins detected in BioID using BMP2K-S as bait.

**Table S3.**
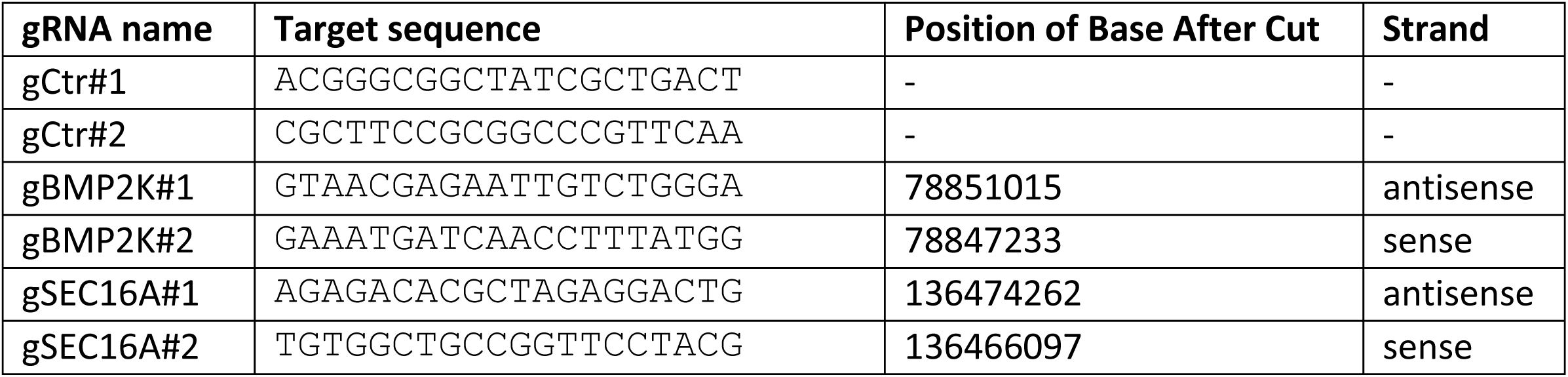
gRNAs used for CRISPR/Cas9 mediated gene inactivation.

**Table S4.**
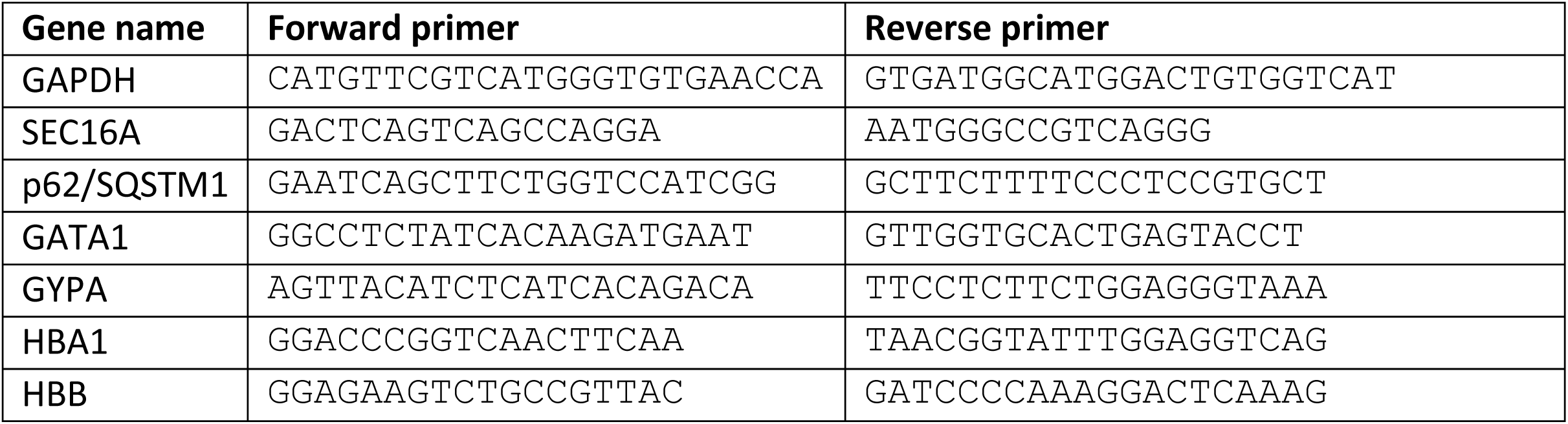
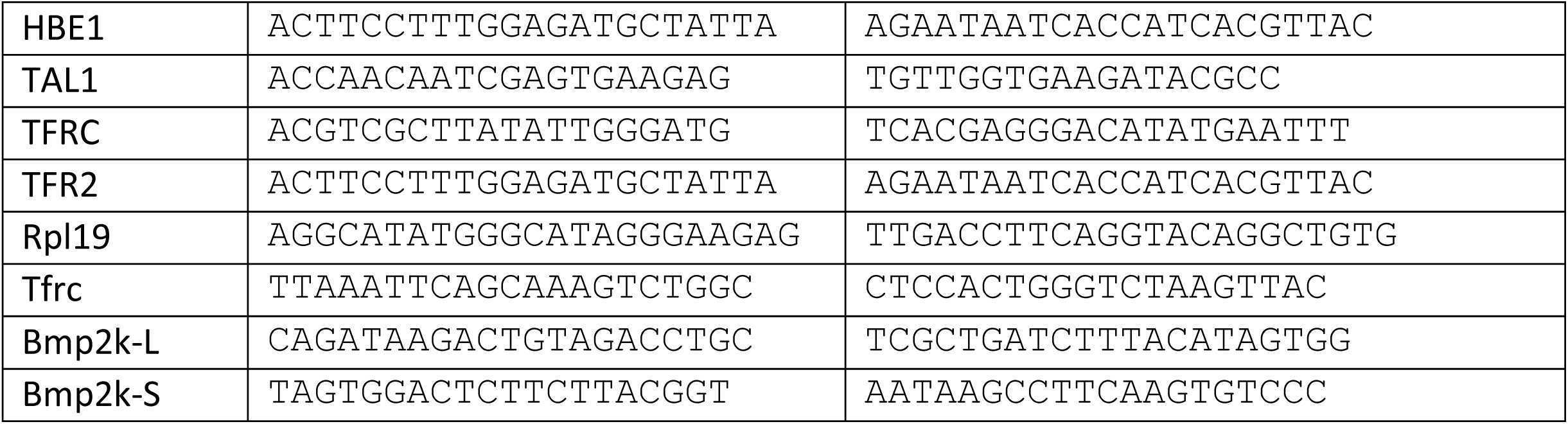
Sequences of primers used for qRT-PCR.

